# Variation in Dental Tissues: Using Bayesian Multilevel Modelling to Explore Intra- and Inter-Individual Dental Variation

**DOI:** 10.1101/2023.05.23.541877

**Authors:** Christianne Fernee, Sonia Zakzewski, Kate Robson Brown

## Abstract

**Objectives:** Dental variation within populations and, even more so, within individuals is far less well understood than variation between populations. This is problematic as a single tooth type is often used as a representative of the whole dentition, despite a lack of understanding of intra-tooth type relationships. This research investigates the variation of dental tissues and proportions within and between individuals.

**Materials and Methods:** Upper and lower first incisor to second premolar tooth rows were obtained from 30 individuals (n=300), from 3 archaeological samples. The teeth were micro-CT scanned and surface area and volumetric measurements were obtained from the surface meshes extracted. Dental variation of these measurements on a tooth and individual level was studied using Bayesian Multilevel Modelling.

**Results:** The individual and tooth level variation differed by dental measurement, ranging between 9.5%-47.5% and 52.6-90.5% respectively. Enamel volume had the highest degree of individual-level variation in contrast to coronal dentine volume that had the lowest of individual-level variation. Tooth type, isomere, and position in field all showed a significant effect on the dental measurements examined in this study.

**Discussion:** Tooth selection and sampling strategies should consider individual and tooth-level variation, with at least one tooth from each type and isomere included in analyses. This will ensure that any population-level differences are not masked by variability between teeth. The low level of coronal dentine volume individual variation indicates that it is particularly useful in studies with small sample sizes.

## 1. Introduction

Data in biological anthropology and osteoarchaeology is inherently nested and identifying variation at the different levels of this data is often at the heart of inquiry. Traditionally, the Bayesian paradigm has been used for chronological modelling. In recent years, however, Bayesian models are being increasingly used to study a wide range of aspects of archaeology (Otárola-Castillo & Torquato 2018). Bayesian multilevel models can be used to answer a variety of questions, regarding both the levels in the respective model and associated predictors. However, Bayesian multilevel modeling is yet to be fully exploited in archaeology and anthropology (Fernée and Trimmis 2021).

Dental variation is multi-layered and multifaceted, occurring within and between individuals and involving the entire tooth or a specific aspect. Variation also occurs over different timescales; over an individuals lifetime and in a population over generations. This diversity is essential for dental performance, responding to external stimuli such as mastication (Brook et al. 2014; Brook et al. 2012). Understanding of such dental variation can give an insight into a number of aspects of human history, including human evolution and migratory patterns (Townsend et al. 2013).

Dental variation within populations and, even more so, within individuals is far less well understood than variation between populations, in both archaeological and clinical research. This is despite, for example, evidence for different developmental programmes for the upper and lower dentitions (Ferguson et al., 2000, 2001; Cobourne and Mitsiadis, 2006; Sperber, 2006) and calls for the need to probe the range and nature of intra-population tooth size variation further (Hemphill 2016a; Hemphill 2016b). This is problematic as a single tooth type is often used as a representative of the whole dentition, despite a lack of understanding of intra-tooth type relationships. Understanding individual level differences, therefore, will have ramifications on between-population studies; How teeth vary between and within individuals may affect whether differences are detected between populations.

Existing intra-population studies, as with inter-population studies, of tooth variation have relied largely on metric crown measurements. Alternative dental measurements, such as tooth tissue proportions, may provide interesting new insights into dental variation. Le Luyer et al (2016; 2018) highlighted this in their research on external and internal phenotypic variation in Pleistocene and Holocene individuals. Like studies of inter-population variation, there is a tendency for intra-population studies to rely on dimensions from a single tooth or tooth class, despite the need for the use of multiple tooth classes (Harris 2003).

The current study investigated inter- and intra-individual variation in dental tissues and proportions using Bayesian Multilevel modelling. The main aim of this research was to explore then main sources of variation in tooth size, between and within individuals.

## 2. Materials and Methods

### 2.1 Materials

The sample studied comprises 300 permanent maxillary and mandibular tooth rows, from first incisors to second premolars, from archaeological individuals (Table 1). Left teeth were primarily analysed, however if unavailable the right tooth was used. The teeth were obtained from 30 individuals from three archaeological samples from the south of the UK. The first sample is derived from 10 individuals from the Early Medieval cemetery at Great Chesterford, Essex (*n=*100), dated from 5^th^ – 7^th^ century AD (Table 1). The second is derived from 10 individuals from the middle Medieval monastic cemetery at Llandough, South Wales (*n=*100) dated from 7^th^ – 11^th^ century AD (Table 1). The final sample was obtained from 10 individuals from the Late Medieval priory cemetery of St Peter and Paul, Taunton, Somerset (*n=*100) dated from 12^th^ – 15^th^ century AD (Table 1).

**Table 1.**
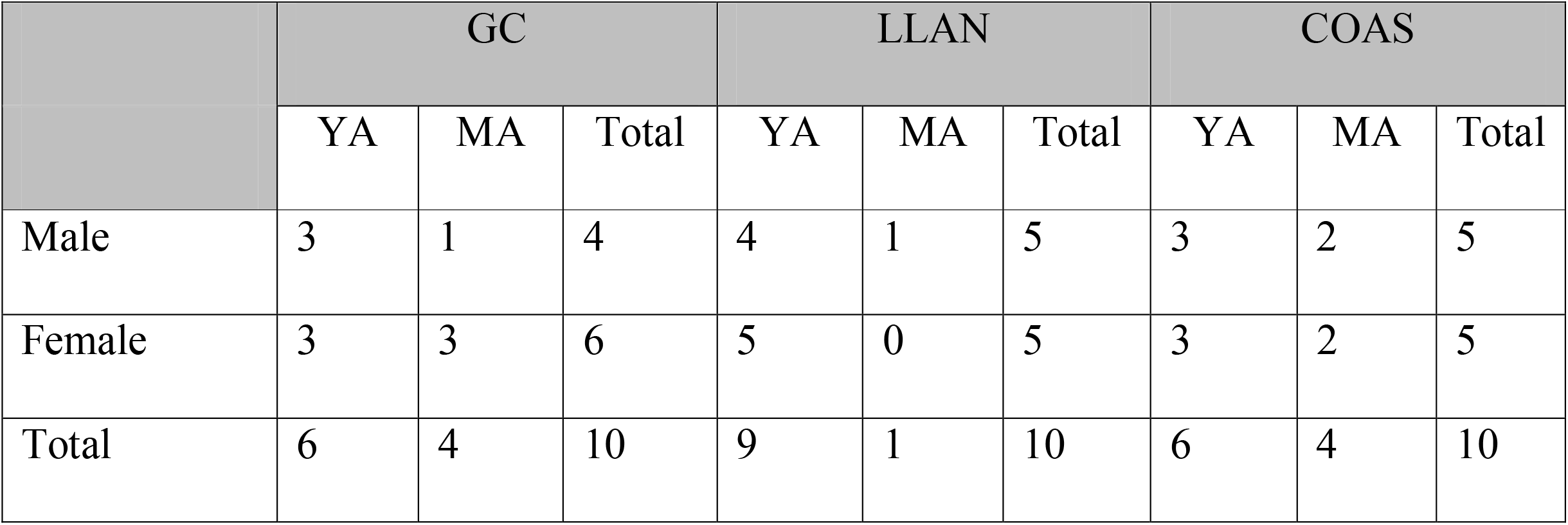
Overall sample composition: distribution of sex and age in each chronological sample. Samples: Great Chesterford (GC), Llandough (LLAN) and Taunton (COAS). Sex: Male and Female. Age categories: Young adult (YA) and Middle adult (MA)

Age and sex were estimated according to established guidelines (Buikstra and Ubelaker, 1994). Sex was estimated based upon the dimorphic characteristics of the pelvis and skull, where available (Buikstra and Ubelaker, 1994, pp 15–21). Age estimates were taken from pelvic characteristics of the pubic symphysis (Todd, 1920; Brooks and Suchey, 1990) and auricular surface (Lovejoy et al., 1985), and were then classified by category: young adult, middle adult and old adult. However, no individuals were classified as an old adult (Table 1). Tooth wear was recorded using Molnar (1971). Individuals with dental anomalies and teeth with pathologies or severe wear (Molnar 1971 >4) were excluded.

### 2.2 Methods: Data Acquisition

Teeth were scanned using micro-computed tomography (micro-CT), employing different scanning facilities and parameters. Loose teeth and teeth in small bony fragments were scanned using a Bruker SkyScan 1272 at the University of Bristol and a Bruker SkyScan 1275 at the Sumitomo Laboratory, Swansea. Loose teeth were scanned at 90 kV and 70 μA using a 0.5 Al & 0.038 Cu filter, for a target resolution of 17.5 μm. Teeth in small bony fragments were scanned at 100 kV and 100 μA using a 1.0 mm Cu filter, for a target resolution of 17.5 μm. Teeth in large bony fragments and crania were scanned using a Nikon XT H 320 at the National Composite Centre (NCC), Bristol, at 145 kV and 110 μA using no filter, for a target resolution of 65 μm (For more details on scan parameters see supporting material 1).

The scans were reconstructed using NRecon 1.7.1.0 (Bruker micro-CT, Belgium) and CTPro3D (Nikon Metrology, Herts UK). The data was then segmented using ScanIP 2018.12 (Simpleware, Exeter, UK) based on thresholding criteria to create individual masks for enamel, dentin, pulp chamber and whole tooth. Cracks were virtually filled in for these masks. For the purposes of this study, cementum was included in the dentine mask as only the external tooth geometry was required. For each threshold range a surface mesh was generated and exported, resulting in four unique meshes for each tooth: enamel, dentin, pulp and whole tooth (Figure 1).

**Figure 1.**
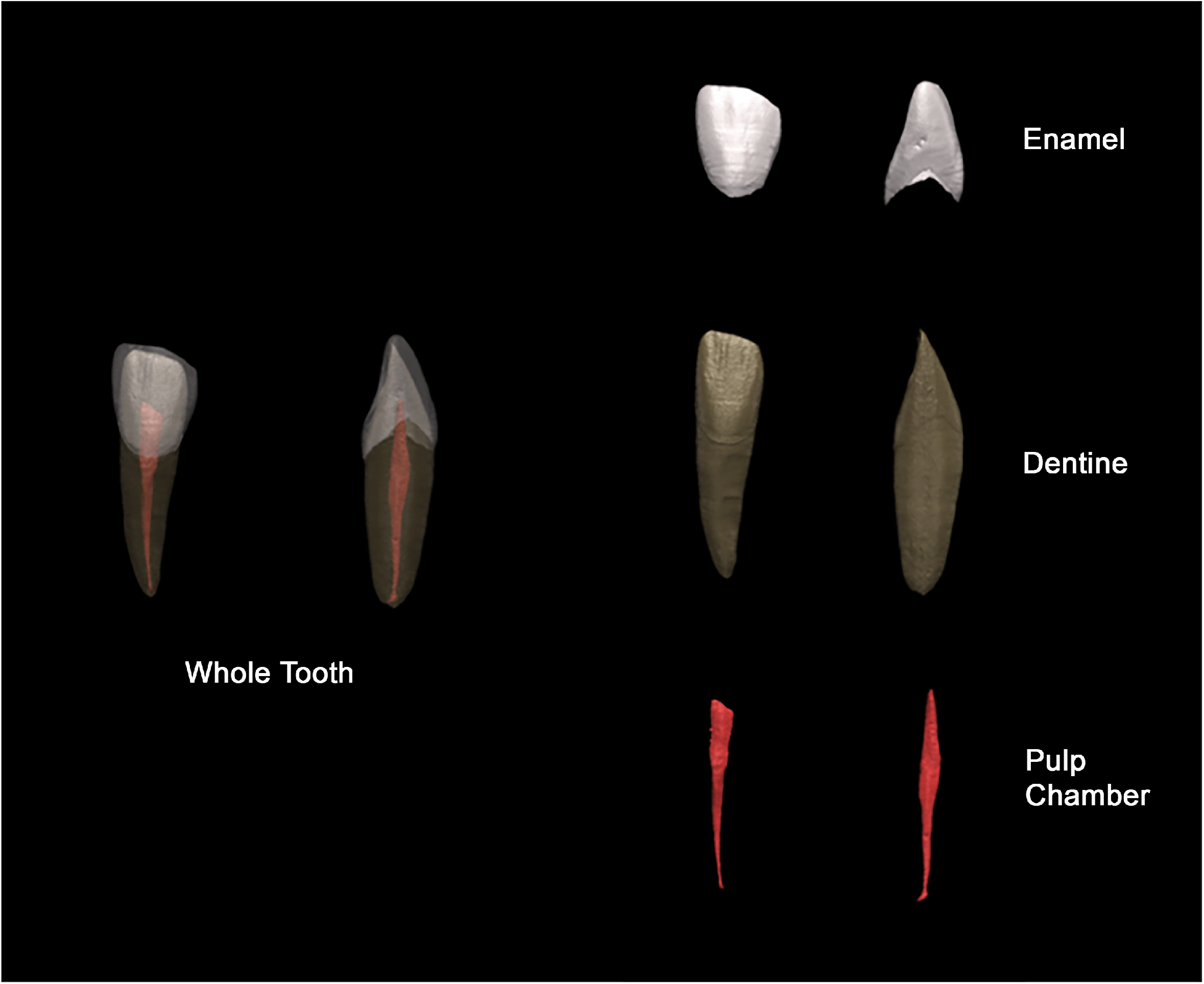
Dental tissue masks from which measurements were taken: whole tooth, enamel, dentine and pulp chamber.

#### 2.2.1 Measurements

Tooth tissue volumes, surface areas and proportions were obtained for each tooth (Table 2; Figure 1 and Figure 2). The root was defined as the tooth present below the CEJ and was extracted in MATLAB R2018a (Mathworks, MA, USA). The surfaces were downsampled prior to separation of the crown and root. Volumetric measurements could only be obtained using closed meshes; meshes were closed in ANSYS ICEM (ANSYS Inc., Canonsburg PA, USA).

**Table 2.**
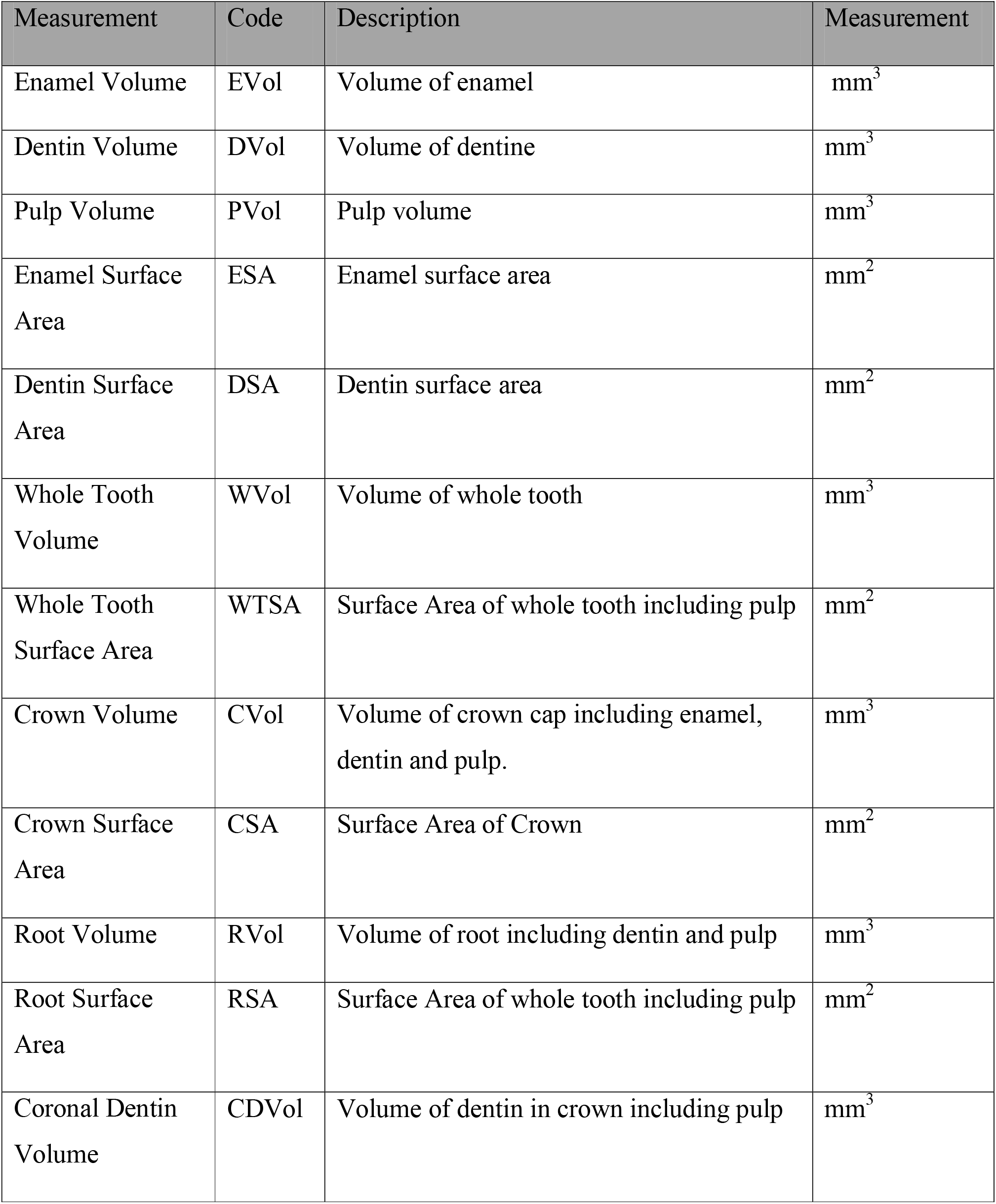
Definition of dental measurements

**Figure 2.**
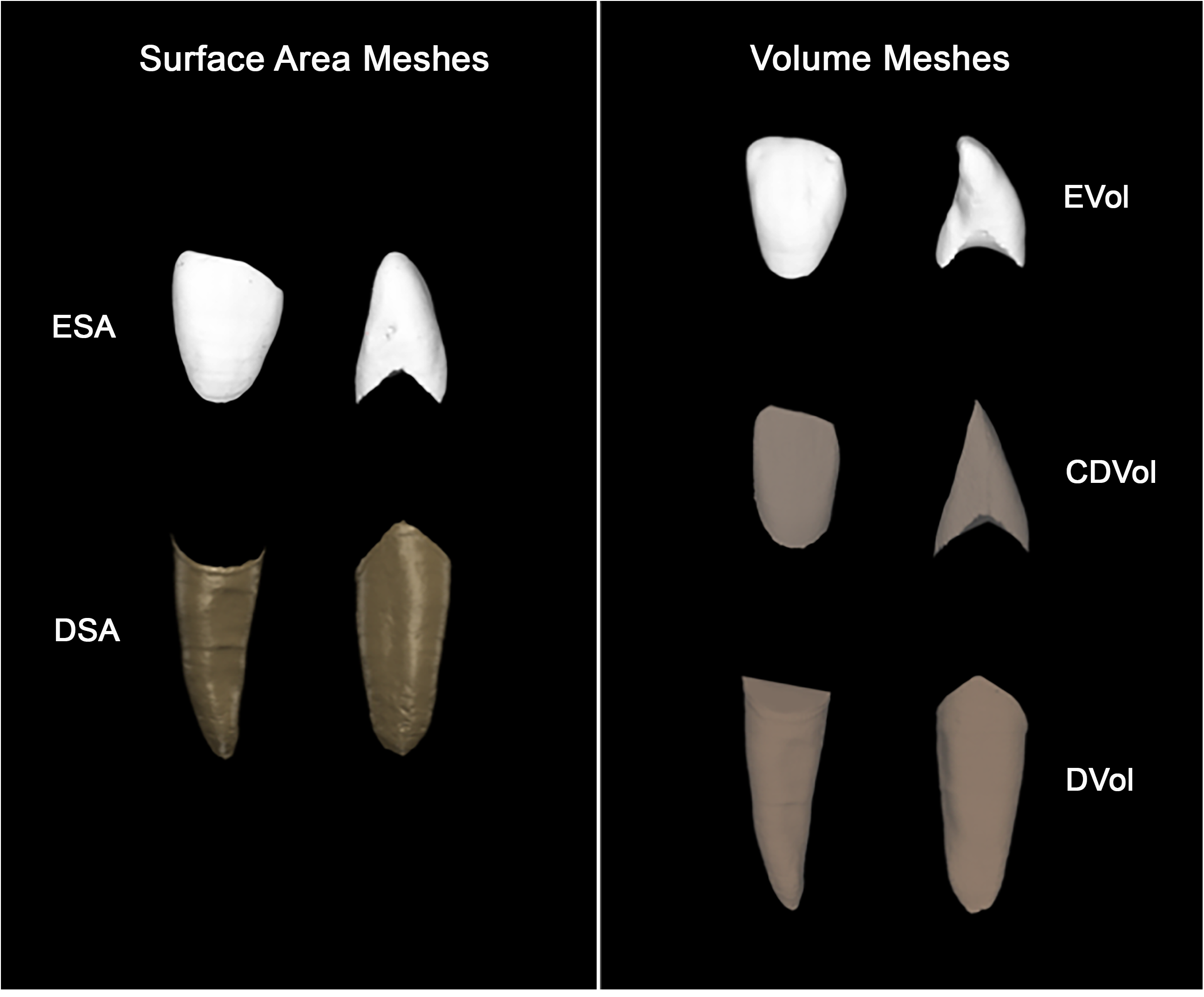
Crown and root surface area and volumetric meshes. Left: Crown surface area (CSA) and root surface area (RSA). Right: Crown volume (CVol), coronal dentine volume (CDVol) and root volume (RVol).

#### 2.2.2 Statistical Analysis: Bayesian multilevel modelling

Dental variation on an intra and inter individual level was studied using Bayesian Multilevel Modelling. Models were constructed and analysed in R 4.1.3 (R Core Team 2021) using the brms package (Bürkner 2017). Posterior distributions were extracted and visualised using the tidybayes package (Kay 2022). The parameters’ uncertainty was summarised by the 95% credible interval (CI).

A Markow Chain Monte Carlo (MCMC) method implemented in Stan (https://mc-stan.org/) with 2 chains of 50000 iterations, with a warm-up of 500, to calculate posterior parameter estimates. Convergence of the chains and autocorrelation was checked visually using trace plots and using convergence diagnostics. These diagnostics include the R-hat, Bulk Effective Sample Size (ESS) and Tail ESS. The R-hat diagnostic provides information on the convergence of the chains and, if chains are not mixed well, will be greater than 1. The ESS corresponds to the number of independent samples. It measures how much independent information there is in autocorrelated chains (Kruschke 2015), with a value of 1,000 sufficient for stable estimates (Bürkner, 2017). The Bulk ESS is a measure for the overall sampling efficiency in the bulk of the distribution, whereas the tail ESS corresponds to the 5% and 95% quantiles. Model assumptions were checked using a QQ-plot.

A stepwise approach was used to construct each model, adding a single parameter at a time. The models were compared using an approximate leave-one-out (LOO) cross-validation, using the loo package (Vehtari et al 2016), where a lower LOO indicates a better model fit and a difference less than 4 interpreted as small (Sivula et al 2020; Bürkner 2017). The LOO procedure chosen as it has numerous advantages over the DIC, which is often used to evaluate the fit of Bayesian models (see Vehtari et al 2017). Changes in the LOO were considered alongside changes in estimates of variance and the posterior mean and standard deviation of the regression (fixed) coefficient evaluate the goodness-of-fit of each model. The MCMC posterior means and 95% credible intervals are presented as summaries of the posterior distribution of each variable.

For each dental measurement a two-level model was constructed: tooth (level 1) and individual (level 2). Specifically, this included 300 teeth, first incisor to second premolar, from 30 individuals. Tooth and individual level predictors were included. At a tooth level: tooth type, isomere, position in field and degree of dental wear were included. At the individual level: sex and age category were included (Figure 3).

**Figure 3.**
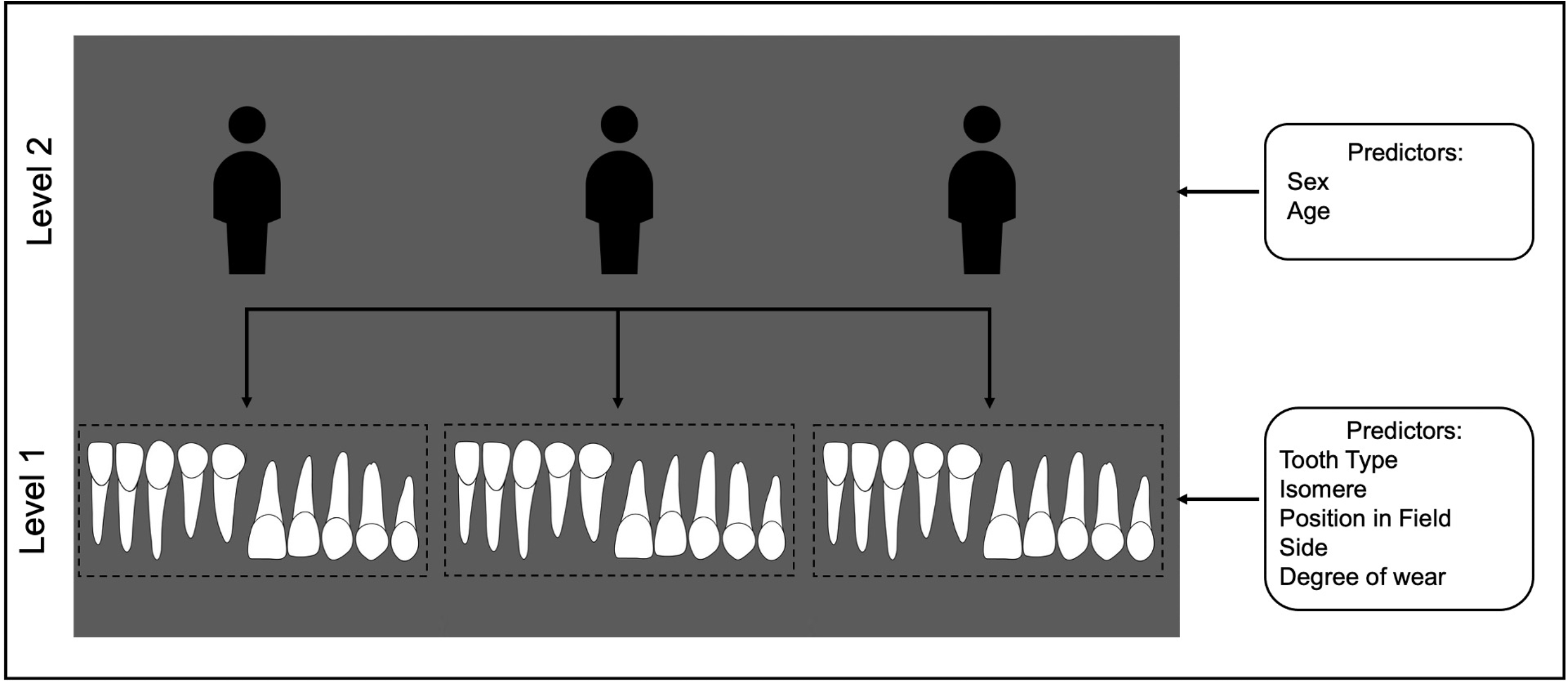
Multilevel Model structure for each multilevel model.

On a tooth level, variation between teeth has been associated with tooth type (Lombardi, 1975; Townsend and Brown, 1979; Harris and Bailit, 1988; Dempsey et al., 1995; Harris, 2003), and between upper and lower dentitions, which are the result of two different developmental programmes (Ferguson et al., 2000, 2001; Cobourne and Mitsiadis, 2006; Sperber, 2006). Position in field was included as a predictor as morphogenetic field theory models suggest that there should be distinct patterns of heritability within each tooth class (Butler 1939; 2001; Dahlberg 1945). According to these models, there are gradients of variation in tooth size and shape from the mesial, or pole tooth, to distal members of each tooth class. The distal later developing tooth, apart from the lateral incisor, shows greater variation (Line, 2001; Townsend et al., 2009a).

On the individual level, variation in dental size has been found by both sex and age. Tooth size has been used in archaeological and modern dentitions to sex individuals using discriminant function analysis (Fernée et al., 2020; Viciano et al., 2011; Tardivo et al., 2015). Tooth size variation by age is largely the product of degenerative processes, with undergoing a range of macroscopic and microscopic degenerative processes, including wear, changes in root chamber morphology, root resorption, changes in root transparency and cementum annulations (Montoya et al., 2015; Schmidt, 2016).

A normal likelihood was fitted to each dental measurement model with an identity link function. Normal distributions were used for each of the random effects (levels) with a weakly informative prior distribution of normal (0, 10), and weakly informative prior of student t (3,0,74.2) was used for the fixed effects (for mathematical structure see supporting information 2).

The data was fitted into 8 models: 1) a null two-level random-intercept model (M_n_), 2) a two-level random-intercept model with one level 2 (L1) predictor (tooth type) (M_1_), 3) a two-level random-intercept model with two L1 predictors (tooth type and isomere) (M_2_), 4) a two-level random-intercept model with three L1 predictors (tooth type, isomere and position in field) (M_3_), 5) a two-level random-intercept model with four L1 predictors (tooth type, isomere, position in field and side) (M_4_), 6) a two-level random-intercept model with five L1 predictors (tooth type, isomere, position in field, side and degree of wear) (M_5_), 7) a two-level random-intercept model with five L1 predictors (tooth type, isomere, position in field, side and degree of wear) and one level 2 (L2) predictor (sex) (M_6_), 8 a two-level random-intercept model with five L1 predictors (tooth type, isomere, position in field, side and degree of wear) and two L2 predictors (sex and age) (M_7_) (Table 3).

**Table 3.** Details of Multilevel models built using stepwise approach with increasing complexity.

| Model Type | Description |
| --- | --- |
| Model 1 ($M_1$) | 2-level model with no fixed explanatory variables. |
| Model 2 ($M_2$) | 2-level model; 1 fixed explanatory variable (tooth type). |
| Model 3 ($M_3$) | 2-level model; 2 fixed explanatory variables (tooth type and isomere). |
| Model 4 ($M_4$) | 2-level model; 3 fixed explanatory variables (tooth type, isomere and position in field). |
| Model 5 ($M_5$) | 2-level model; 3 fixed explanatory variables (tooth type, isomere, position in field and side). |
| Model 6 ($M_6$) | 2-level model; 5 fixed explanatory variables (tooth type, isomere, position in field, side and degree of wear). |
| Model 7 ($M_7$) | 2-level model; 5 fixed explanatory variables (tooth type, isomere, position in field, side, degree of wear and sex). |
| Model 8 ($M_8$) | 2-level model; 4 fixed explanatory variables (tooth type, isomere, position in field, sex and age). |

#### 2.3.3 Variable Analysis

The proportion of residual variance attributed to each level of the model can be determined by calculating the Variance Partition Coefficient (VPC). The VPC, otherwise known as the intraclass correlation (ICC), for a model fit to continuous data is equal to the intra-unit correlation which is the correlation between level 1 units in the same level 2 unit (Rasbash et al. 2017). For example, teeth within the same dentition. For a two-level model the VPC is calculated by dividing the level 2 residual variance by the combined level 2 and level 1 residual variance. For each model, the proportion of variance of each level was calculated using the performance package (Lüdecke et al. 2021), this was then converted into a percentage to estimate the percentage of residual variance at each level.

The effect of the fixed predictor variables on the outcome variable was interrogated using the posterior distributions, particularly the credible intervals. If the credibility intervals are strictly positive or negative this indicates a significant effect. However, if the interval encompasses 0 the effect appears to be non-significant. In addition to this, the narrower the interval the more precise the estimate effect.

## 3. Results

The visual inspection of the trace plots and the MCMC diagnostics indicated Markov Chain convergence and the normality plots indicated that the residuals at each level were roughly normally distributed (see supporting information 3 and 4 for diagnostic statistics, diagnostic plots and normality plots).

### 3.1 Model Fit Overview

Following the evaluation of the goodness-of-fit of each model, the final models for each dental measurement determined to be M_7._ The M7 LOO was lowest for most models. For the other models there was only a small difference between M7 LOO and the lowest LOO value (see supporting information 5).

### 3.2 Models

#### 3.2.1 Dental Tissues

##### 3.2.1.1 Enamel Surface Area

Variation in the ESA MLM was 58.6% at the tooth level and 41.4% at the individual level (Table 4). For the fixed effects, tooth type (I< C<PM), isomere (L<U), position in field (D<P) and degree of wear (1 < 4) had a significantly affected ESA (Figure 4; Supporting information 6.1). The remaining predictors did not have a significant effect on ESA, with side having a very negligible effect on ESA (Figure 4).

**Table 4.** Summary of multilevel model results for each mode

|  | VPC |  |
| --- | --- | --- |
|  | Level 1 | Level 2 |
| ESA | 58.6% | 41.4% |
| DSA | 58.9% | 41.1% |
| EVol | 52.6% | 47.5% |
| DVol | 75.8% | 24.2% |
| PVol | 63% | 37% |
| CSA | 73.1% | 26.9% |
| RSA | 70.0% | 30.0% |
| CVol | 73.8% | 26.2% |
| RVol | 73.8% | 26.2% |
| CDVol | 90.5% | 9.5% |
| WTSA | 61.0% | 39.0% |
| WTVol | 81.4% | 18.6% |

**Figure 4.**
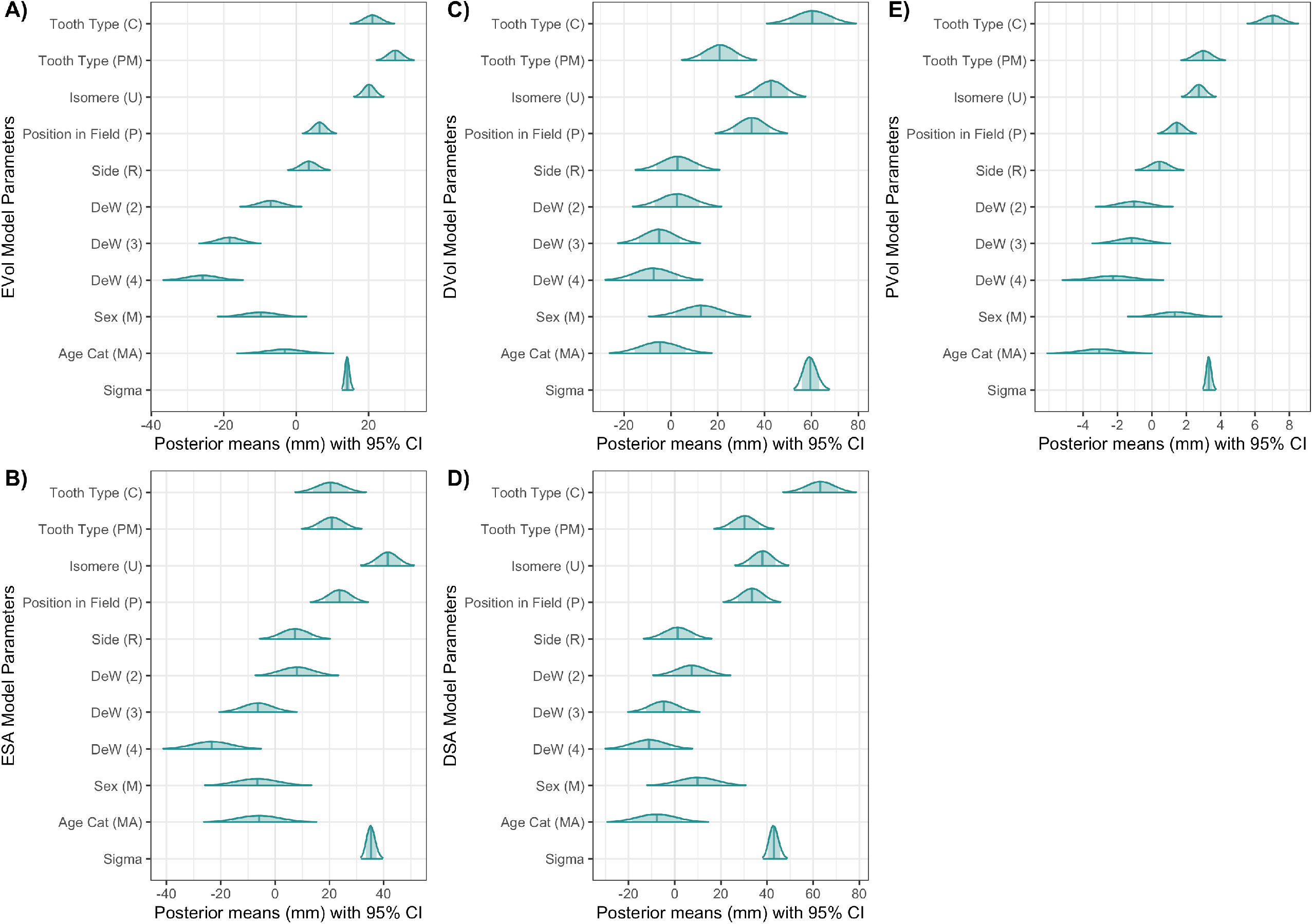
Multilevel model parameter posterior means and 95% credible intervals (CIs) for: A) Enamel Volume (EVol), B) Enamel Surface Area (ESA), C) Dentin Volume (DVol), D) Dentin Surface Area (DSA) and E) Pulp Volume (PVol)

##### 3.2.1.2 Dentin Surface Area

Variation in the DSA MLM was 58.9% at the tooth level and 41.1% at the individual level (Table 4). Tooth type (I<PM<C), isomere (L<U) and position in field (D<P) had significantly affected DSA (Figure 4). Conversely, DSA was not significantly affected by the remaining predictors, especially side (Figure 4).

##### 3.2.1.3 Enamel Volume

Variation in the EVol MLM was 52.5% at the tooth level and 47.5% at the individual level (Table 4). Tooth type (I<C<PM), isomere (L<U), position in field (D<P) and degree of wear (1>3, 1>4) significantly affect EVol (Figure 4). Conversely, the remaining predictors did not significantly affect EVol, with a very negligible effect of side EVol (Figure 4).

##### 3.2.1.4 Dentin Volume

Variation in the DVol MLM was 75.8% at the tooth level and 24.2% at the individual level (Table 4). Tooth type (I<PM<C), isomere (L<U) and position in field (D<P) significantly affected DVol (Figure 4). Conversely, the remaining predictors did not significantly affect DVol, again side had a very negligible effect on DVol (Figure 4).

##### 3.2.1.5 Pulp Volume

Variation in the PVol MLM was 52.5% at the tooth level and 47.5% at the individual level (Table 4). Tooth type (I<C<PM), isomere (L<U), position in field (D<P), degree of wear (1>4) and age (YA > MA) significantly affected PVol (Figure 4). Conversely, the remaining predictors did not significantly affect PVol (Figure 4).

#### 3.2.2 Tissue Proportions

##### 3.2.2.1 Crown Surface Area

##### 3.2.2.2

Variation in the CSA MLM was 73.1% at the tooth level and 26.9% at the individual level (Table 4). Tooth type (I<C<PM), isomere (L<U), position in field (D<P) and degree of wear (1>4) significantly affected CSA (Figure 5). The remaining predictors did not significantly affect CSA, with side having a very negligible effect (Figure 5).

**Figure 5.**
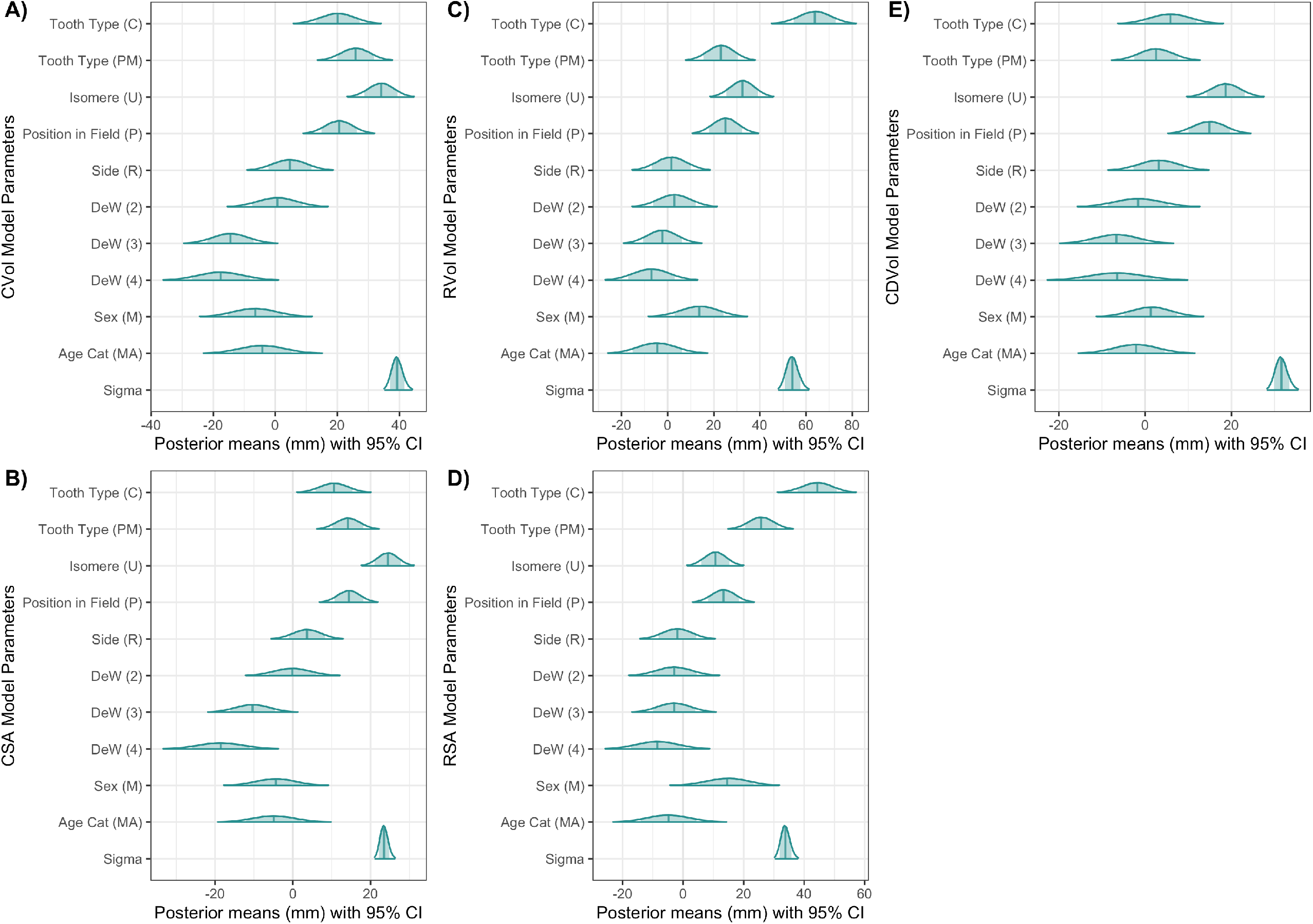
Multilevel model parameter posterior means and 95% credible intervals (CIs) for: A) Crown Volume (CVol), B) Crown Surface Area (CSA), C) Root Volume (RVol), D) Root Surface Area (RSA) and E) Coronal Dentin Volume (CDVol)

##### 3.2.2.1 Root Surface Area

Variation in the RSA MLM was 70.0% at the tooth level and 30.0% at the individual level (Table 4). Tooth type (I<PM<C), isomere (L<U) and position in field (D<P) significantly affected RSA (Figure 5). The remaining predictors did not significantly affect RSA (Figure 5).

##### 3.2.2.1 Crown Volume

Variation in the CVol MLM was 73.8% at the tooth level and 26.2% at the individual level (Table 4). Tooth type (I<PM<C), isomere (L<U), position in field (D<P) and degree of wear (1>3, 1>4) significant affected CVol (Figure 5). The remaining predictors did not significantly affect CVol (Figure 5).

##### 3.2.2.1 Root Volume

Variation in the RVol MLM was 73.8% at the tooth level and 26.2% at the individual level (Table 4). Tooth type (I<PM<C), isomere (L<U) and position in field (D<P) significantly affected RVol (Figure 5). The remaining predictors did not significantly affect RVol (Figure).

##### 3.2.2.1 Coronal Dentin Volume

Variation in the CDVol MLM was 90.5% at the tooth level and 9.5% at the individual level (Table 4). Tooth type (I<PM<C), isomere (L<U), position in field (D<P) and age (YA<MA) significantly affected CDVol (Figure 5). The remaining predictors did not significantly affect CDVol (Figure 5).

#### 3.2.3 Whole Tooth

##### 3.2.3.1 Whole Tooth Surface Area

Variation in the WTSA MLM was 61.0% at the tooth level and 39.0% at the individual level (Table 4). Tooth type (I<PM<C), isomere (L<U) and position in field (D<P) significantly affected WTSA (Figure 6). The remaining predictors did not significantly affect WTSA (Figure 6).

**Figure 6.**
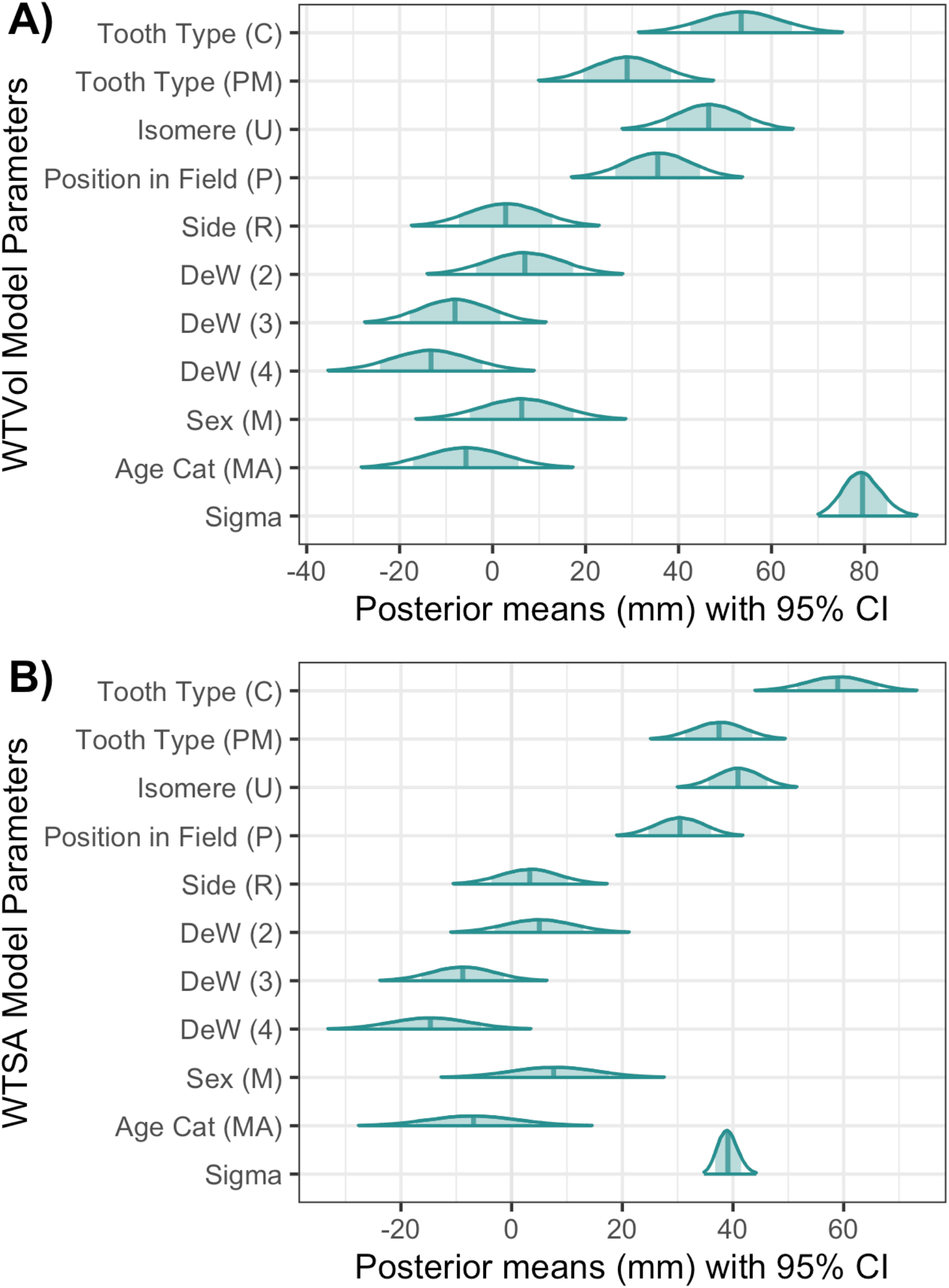
Multilevel model parameter posterior means and 95% credible intervals (CIs) for: A) Whole Tooth Volume (WTVol) and B) Whole Tooth Surface Area (WTSA).

##### 3.2.3.1 Whole Tooth Volume

Variation in the WTVol MLM was 81.4% at the tooth level and 18.6% at the individual level (Table 4). Tooth type (I<PM<C), isomere (L<U) and position in field (D<P) significantly affect WTVol (Figure 6). Conversely, the remaining predictors did not significantly affect WTVol, with side having a very negligible effect on WTVol (Figure 6).

## 4. Discussion

### 4.1 Random Effects

Little is known about intra-dentition variation and calls have been made for research into variation within populations and individuals (Kieser, 1990; Harris, 2003, p 88; Hemphill, 2016). The individual and tooth level variation differed by dental measurement, ranging between 9.5%-47.5% and 52.6-90.5% respectively. However, most measurements averaged around 30% at the individual and 70% at the tooth level. The MLM results indicate that even though a greater proportion of variation occurs between teeth, there remains a considerable amount of variation at an individual level. This has important repercussions on past, present and future research in terms of sample composition. The use of a small number or single individual to represent a whole population or even species may mask the true variation occurring between the compared populations or species.

The degree of variability at the tooth level, although greater than at the individual level in most measurements, is lower than expected. These findings suggest that teeth within the same individual exhibit a considerable degree of similarity, providing justification for the use of dimensions from a single tooth or tooth class in inter-population studies. However, tooth type was found to contribute the most variation to the models for most dental measurements, followed by tooth isomere. Consequently, it is recommended that when possible, at least one tooth from each tooth type and isomere should be included in analyses. This will ensure that the full range of variation within the dentition is captured, and that any population-level differences are not masked by the inherent variability of individual teeth. Overall, these results have important implications for future dental studies, emphasising the need for careful consideration of tooth selection and sampling strategies in analyses of dental variation.

Enamel volume was found to have the highest degree of individual-level variation, accounting for 47.5% of residual variation in the final model. This high degree of variation at the individual level suggests that detecting true between-sample differences in enamel volume may be more difficult than for other dental measurements examined in the study. Consequently, studies that rely solely on enamel volume as a metric may be limited in their ability to accurately characterise population-level variation. It is important for researchers to carefully consider the sources of individual variation in enamel volume, such as age, sex, and diet, and to account for these factors when conducting comparative studies.

In contrast to the other dental measurements examined in this study, coronal dentine volume displayed the least variation between individuals, with only 9.5% of residual variation attributed to the individual level in the final model. This low level of individual variation indicates that coronal dentine volume may be particularly useful in studies with small sample sizes, including studies involving extinct fossil species. Indeed, past paleoanthropological studies have already demonstrated the utility of coronal dentine volume in this regard (Bayle et al., 2017; Buti et al., 2017). The reliability of this measurement may be due to its limited sensitivity to factors that influence other dental measurements, such as dental wear and enamel thickness. Consequently, coronal dentine volume may serve as a valuable metric in studies of dental morphology and evolution, particularly when sample sizes are limited.

### 4.2 Fixed Effects

Tooth type, isomere, and position in field all showed a significant effect on the dental measurements examined in this study. These findings are in line with existing knowledge about the sources of variation in dental morphology and their developmental origins (Ferguson et al., 2000, 2001; Cobourne and Mitsiadis, 2006; Sperber, 2006). As a result, it is recommended that future studies take these factors into account when selecting teeth for analysis and interpreting results. The significance of these variables underscores the importance of understanding the complex interplay between genetic and environmental factors in dental development and evolution.

The study did not find any significant effect of sex or side on the dental measurement models under investigation. It is possible that the lack of significant effect of sex on dental measurements is due to sexual dimorphism being more pronounced in certain types of teeth, such as canines, as suggested by previous studies (Fernée et al., 2020; García-Campos et al., 2018a; b). The negligible effect of side on the measurements is consistent with previous evidence from linear measurements, which indicates that dental asymmetry is minimal and does not affect comparisons (Hillson, 2005). The small number of right-sided teeth included in the sample may have contributed to this result, as the left side was preferentially selected and the right was only included if the left side was unavailable or did not meet the inclusion criteria.

### 4.3 Method

Multilevel modelling presents a powerful tool for addressing a variety of research questions in biological anthropology. By incorporating both fixed and random effects, the complexity of the model can be increased, allowing for the exploration of problematic questions across various levels and associated predictors. The versatility of MLMs extends beyond continuous data, including linear measurements, geometric morphometric co-ordinates, and stable isotope data, to also include categorical, proportion, and ordinal data, as well as time-series and geospatial data. Fernée and Trimmis (2021) have highlighted the broad applicability of MLMs to diverse datasets. Moreover, MLMs are user-friendly, easily accessible, and can be implemented in R and a range of other freely available software packages, such as those discussed by Mai and Zang (2018). Therefore, the potential of MLMs to contribute to the advancement of biological anthropology research is undeniable.

## Conclusion

This study sheds light on the sources and magnitude of variation in dental morphology, highlighting the importance of careful tooth selection and sampling strategies in analyses of dental variation. The results suggest that tooth type and isomere play a significant role in dental measurements, while sex and side have negligible effects. Enamel volume displays the highest degree of individual-level variation, indicating that studies relying solely on this metric may be limited in their ability to accurately characterise population-level variation. On the other hand, coronal dentine volume exhibits the least variation between individuals, making it a valuable metric in studies with small sample sizes. Multilevel modelling is a powerful tool for exploring complex research questions across various levels and associated predictors, with broad applicability to diverse datasets. Overall, these findings have important implications for future dental studies, emphasising the need for careful consideration of tooth selection and sampling strategies in analyses of dental variation.

## Acknowledgements

The data for this paper was obtained during the PhD research of C.L.F which was funded by AHRC-SWW DTP and BABAO.

## Data Availability Statement

The data and code that was used for the analyses is freely available at the following link: https://figshare.com/s/d6a3c1e7125efc3968ac.

## Notes

### Competing Interest Statement

The authors have declared no competing interest.

