## supporting information for "Variation in Dental Tissues: Using Bayesian Multilevel Modelling to Explore Intra- and Inter-Individual Dental Variation"

### 1 Supporting Information: Scan Parameters

| Type of sample | Scan Location | Hardware | Voxel size (µm) | kVp / µA | Pixels | Filter | Number of proj | Frames per proj | Exposure Time (ms) | Source-to-Object distance (mm) | Source-to-Detect distance (mm) | Rotation ($^0$) | Rotation step ($^0$) |
|---|---|---|---|---|---|---|---|---|---|---|---|---|---|
| Loose Teeth | Imaging Lab - University of Bristol | Skyscan 1272 | 17.5 | 90 / 70 | 1224 x 820 | 0.5 Al & 0.038 Cu | 300 | 3 | ~800 | 101.75 | 172.29 | 180 | 0.60 |
| Small bony fragments | Imaging Lab - University of Bristol | Skyscan 1272 | 17.5 | 100 / 100 | 1224 x 820 | 1.0 mm Cu | 300 | 3 | ~900 | 132.715 | 224.72 | 180 | 0.60 |
| | Lab – Sumitomo Swansea | Skyscan 1275 | 17.5 | 100 / 100 | 1944 x 1536 | 1.0 mm Cu | 900 | 6 | ~ 300 | 61.730 | 286.0 | 180 | 0.20 |
| Large bony fragments | NCC | Nikon XT H 320 | 65 | 145 / 110 | 1242 x 827 | None | 2000 | 4 | 500 | 309.54 | 1005.16 | 180 | 0.18 |

### 2 Supporting Information: Additional Model Information

A normal likelihood was fitted to continuous response data (dental measurement) with an identity link function. Normal distributions were used for each of the random effects (levels: tooth and individual) with a student t (3, 0, 74.2) prior distribution. For the fixed parameters (tooth type, isomere, position in field, Side, Degree of wear, Sex and Age) a weakly informative prior of normal (0, 10) was used. A two-level random-intercept model was constructed, the mathematical structure of the final model is as follows:

$$y_{ij} = \beta_{0ij} + \beta_1 X_{1ij} + \beta_2 X_{1ij} + \beta_3 X_{1ij} + \beta_4 X_{1ij} + \beta_5 X_{1ij} + \beta_6 X_{1ij} + \beta_7 X_{1ij}$$
$$\beta_{0ij} = \beta_0 + v_{0k} + u_{0j} + e_{ij}$$

$$u_{0jk} \sim \mathrm{N}(0, \sigma_{u0}^2)$$
$$e_{ijk} \sim \mathrm{N}(0, \sigma_e^2)$$

Prior Specifications

p($\beta_0$) ~ normal (0,10)
p($\beta_1$)~ normal (0,10)
p($\beta_2$)~ normal (0,10)
p($\beta_3$)~ normal (0,10)
p($\beta_4$)~ normal (0,10)
p($\beta_5$)~ normal (0,10)
p($\beta_6$)~ normal (0,10)
p($\beta_7$)~ normal (0,10)
p($1/\sigma_{u0}^2$) ~ student_t(3, 0, 74.2)
p($1/\sigma_e^2$)~ student_t(3, 0, 74.2)

where $y_{ijk}$ is the dental measurement (mm) ($y$) of each tooth $i$ within an individual $j$. $\beta_0$, $\beta_1 X_{1ij}$, $\beta_2 X_{1ij}$, $\beta_3 X_{1ij}$, $\beta_3 X_{1ij}$, $\beta_4 X_{1ij}$, $\beta_5 X_{1ij}$, $\beta_6 X_{1ij}$ and $\beta_7 X_{1ij}$ is the fixed part of the model, $v_{0k} + u_{0jk}$ is the random part of the model and $\beta_{0ij}$ is the overall error term.

### 3 Supporting Information: Checking Model Assumptions

#### 3.1 Tissue Volume and Surface Area Model QQ Plots

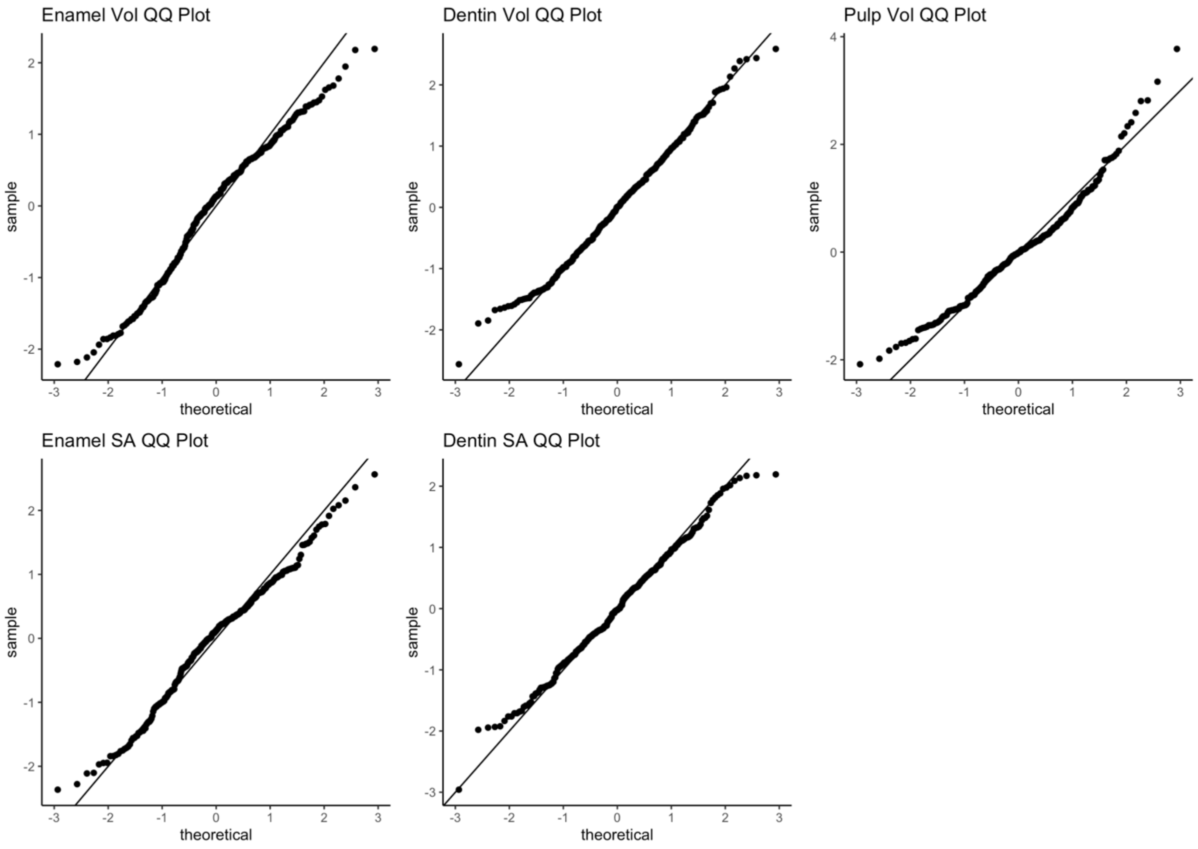

#### 3.2 Dental Proportion Volume and Surface Area Model QQ Plots

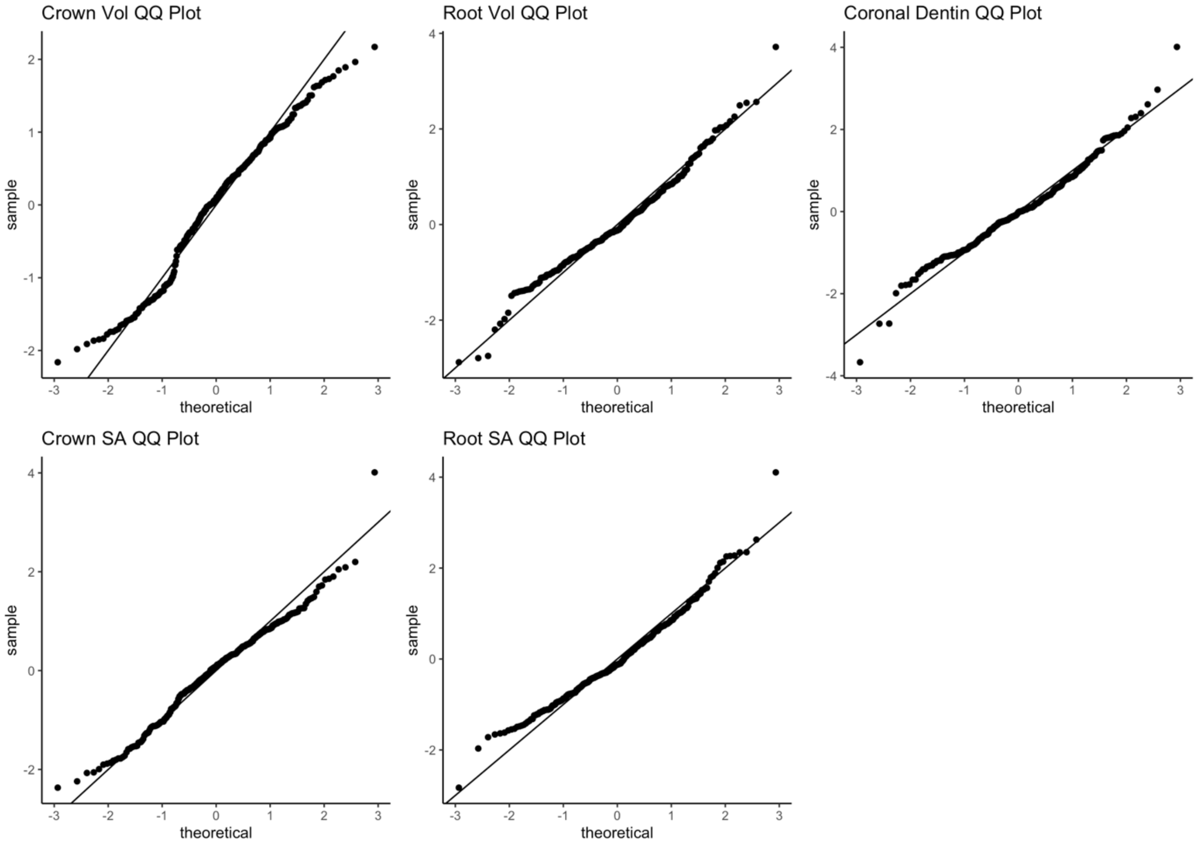

#### 3.3 Whole Tooth Volume and Surface Area Model QQ Plots

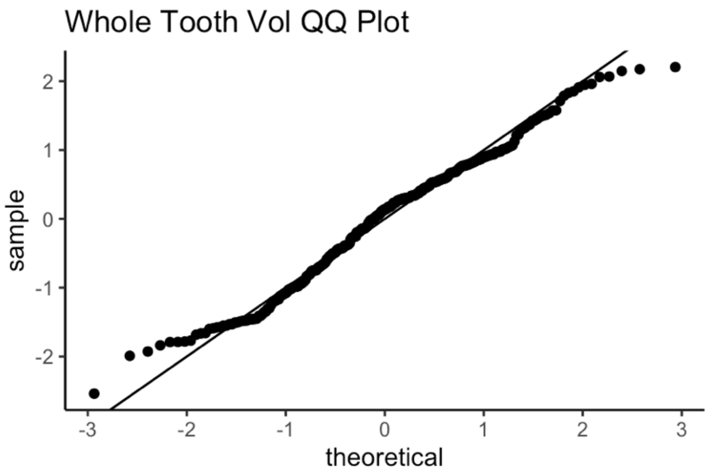

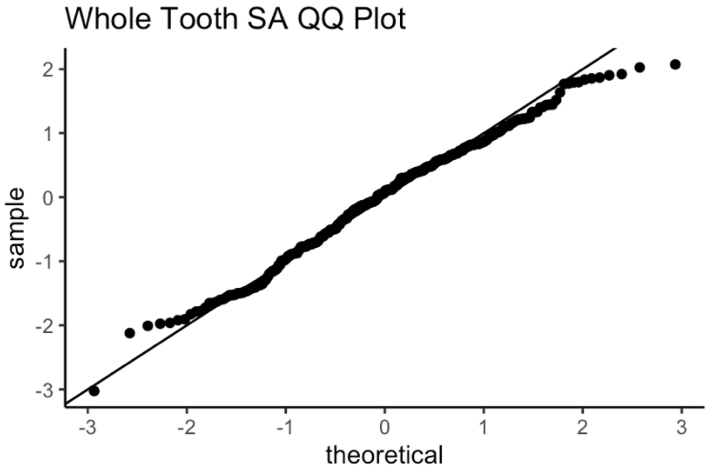

### 4 Supporting Information: Model Convergence Checks

#### 4.1 Enamel Volume

*Table 1 MCMC diagnostics for the Enamel Volume Model*

| Parameter | Rhat | Bulk ESS | Tail ESS |
|---|---|---|---|
| Random | | | |
| Individual | 1.00 | 24726 | 44201 |
| Tooth | 1.00 | 105847 | 75076 |
| Fixed | | | |
| Intercept | 1.00 | 28006 | 48956 |
| Tooth Type (C) | 1.00 | 110760 | 81378 |
| Tooth Type (PM) | 1.00 | 77645 | 77503 |
| Isomere (U) | 1.00 | 154200 | 74003 |
| Position in Field (P) | 1.00 | 144961 | 79607 |
| Side (R) | 1.00 | 117997 | 77355 |
| DeW 2 | 1.00 | 55944 | 70482 |
| DeW 3 | 1.00 | 45735 | 64197 |
| DeW 4 | 1.00 | 48206 | 66862 |
| Sex (M) | 1.00 | 24584 | 43839 |
| Age (MA) | 1.00 | 26965 | 47228 |

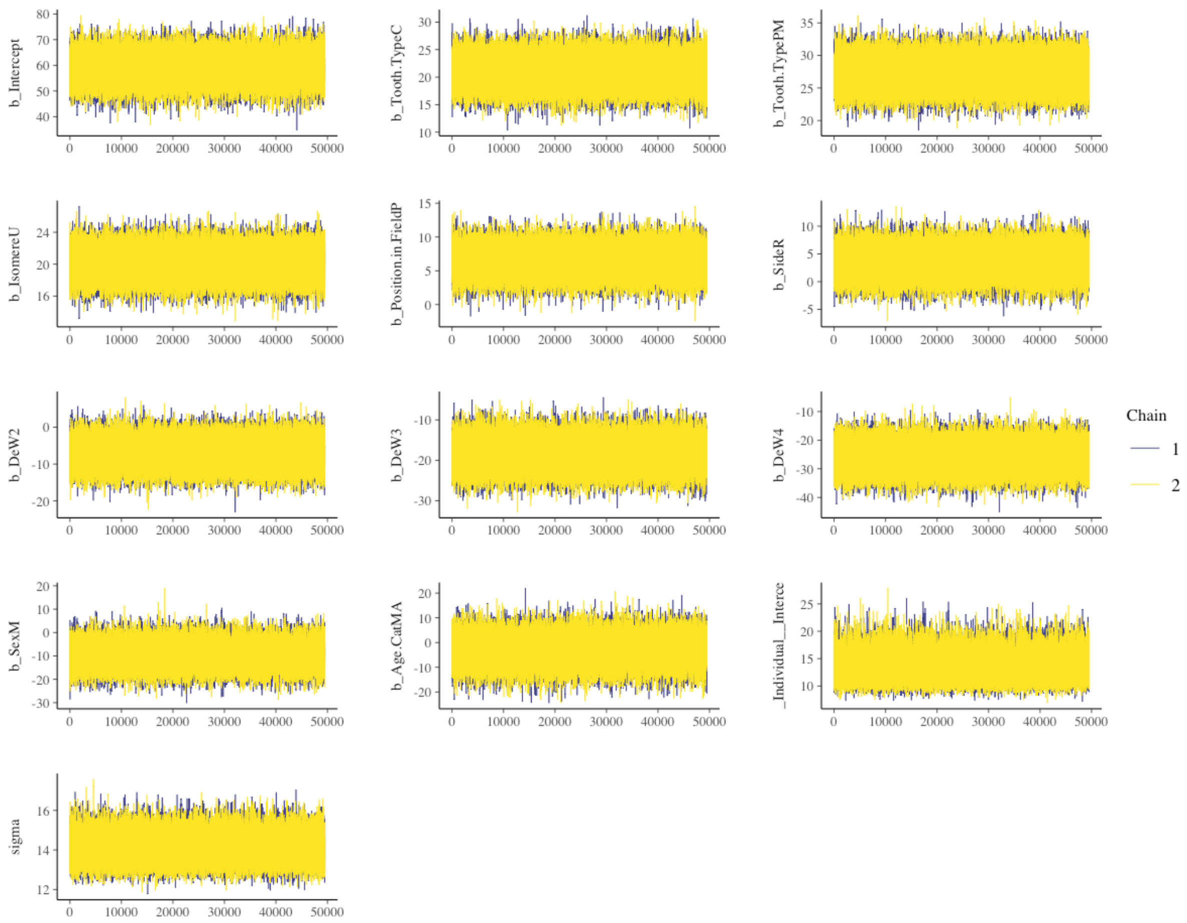

*Figure 1 Enamel volume M7 model Markov Chain Monte Carlo Trace for each parameter.*

#### 4.2 Enamel Surface Area

*Table 2 MCMC diagnostics for the Enamel Surface Area Model*

| Parameter | Rhat | Bulk ESS | Tail ESS |
|---|---|---|---|
| Random | | | |
| Individual | 1.00 | 32017 | 47299 |
| Tooth | 1.00 | 122284 | 77021 |
| Fixed | | | |
| Intercept | 1.00 | 50454 | 64933 |
| Tooth Type (C) | 1.00 | 140327 | 81488 |
| Tooth Type (PM) | 1.00 | 133053 | 77876 |
| Isomere (U) | 1.00 | 181592 | 74096 |
| Position in Field (P) | 1.00 | 171006 | 78548 |
| Side (R) | 1.00 | 175375 | 74340 |
| DeW 2 | 1.00 | 124415 | 84154 |
| DeW 3 | 1.00 | 104806 | 81672 |
| DeW 4 | 1.00 | 103845 | 80825 |
| Sex (M) | 1.00 | 60530 | 70236 |
| Age (MA) | 1.00 | 66668 | 71916 |

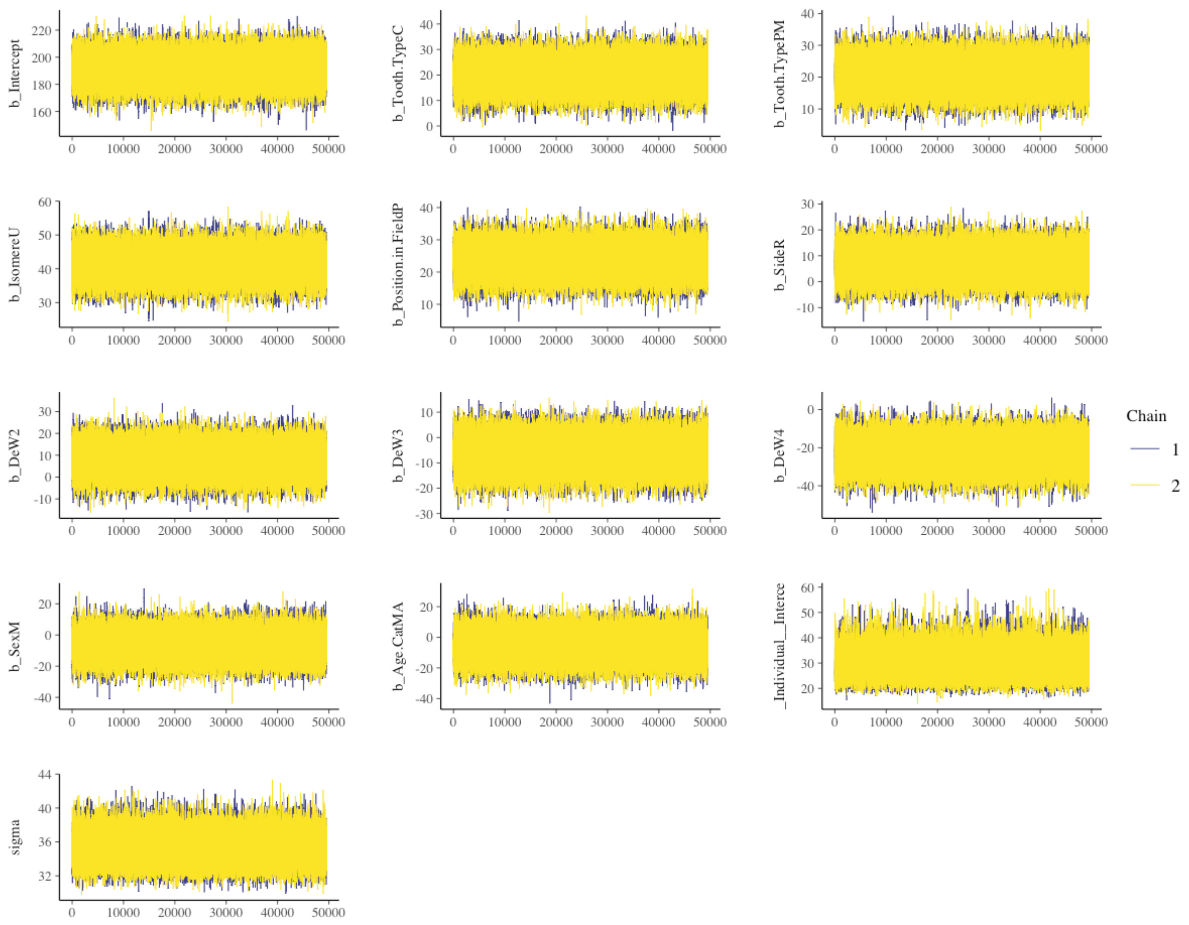

*Figure 2 Enamel surface area M7 model Markov Chain Monte Carlo Trace for each parameter*

#### 4.3 Dentin Surface Area

*Table 3 MCMC diagnostics for the Dentin Surface Area Model*

| Parameter<br>Random | Rhat | Bulk ESS | Tail ESS |
|---|---|---|---|
| Individual | 1.00 | 32413 | 53566 |
| Tooth | 1.00 | 113142 | 78211 |
| Fixed | | | |
| Intercept | 1.00 | 42098 | 62512 |
| Tooth Type (C) | 1.00 | 137076 | 79880 |
| Tooth Type (PM) | 1.00 | 148929 | 76828 |
| Isomere (U) | 1.00 | 190008 | 74064 |
| Position in Field (P) | 1.00 | 151426 | 76988 |
| Side (R) | 1.00 | 176315 | 76469 |
| DeW 2 | 1.00 | 114845 | 82251 |
| DeW 3 | 1.00 | 100717 | 80265 |
| DeW 4 | 1.00 | 101293 | 80465 |
| Sex (M) | 1.00 | 60281 | 73908 |
| Age (MA) | 1.00 | 71605 | 77509 |

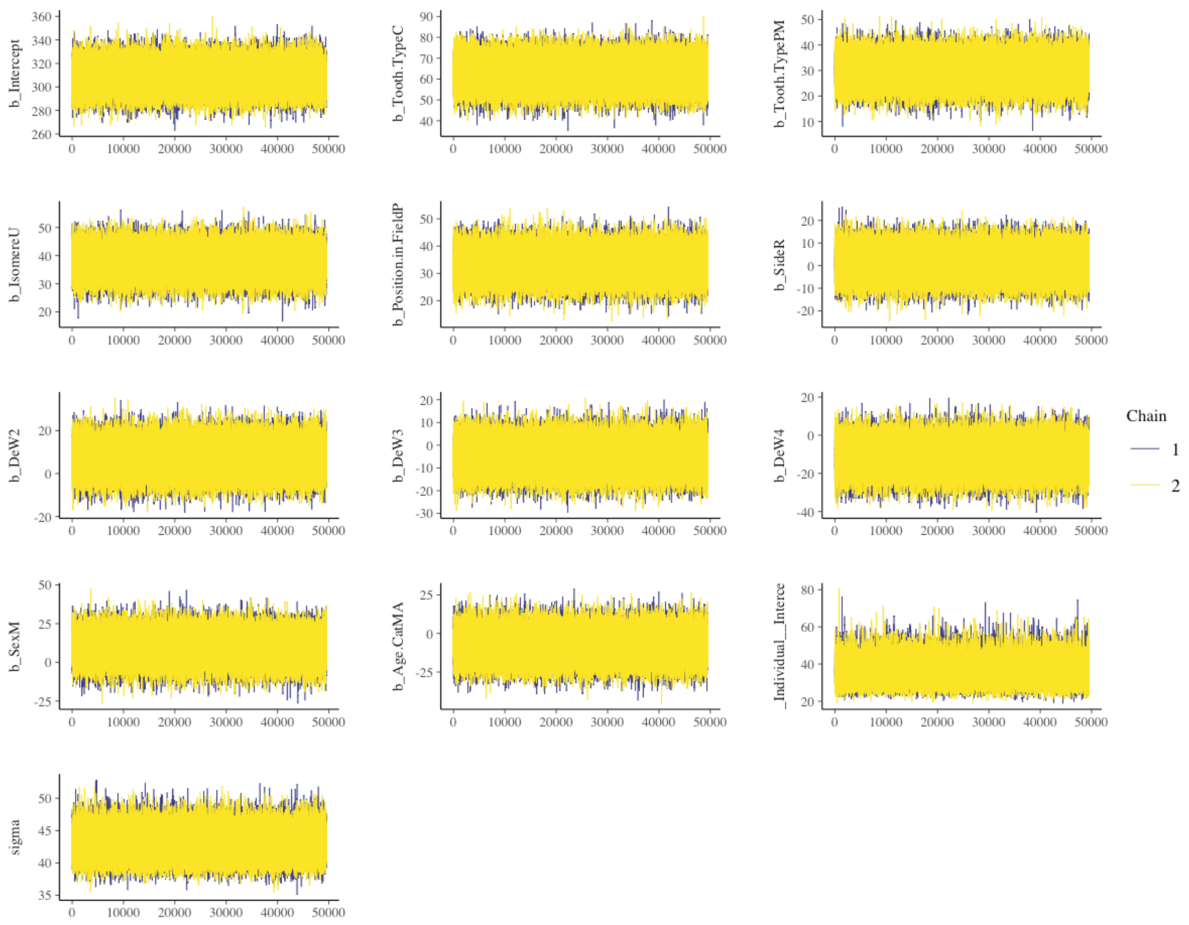

*Figure 3 Dentin surface area M7 model Markov Chain Monte Carlo Trace for each parameter.*

#### 4.4 Dentin Volume

*Table 4 MCMC diagnostics for the Dentin Volume Model*

| Parameter | Rhat | Bulk ESS | Tail ESS |
|---|---|---|---|
| Random | | | |
| Individual | 1.00 | 37947 | 55532 |
| Tooth | 1.00 | 105356 | 77729 |
| Fixed | | | |
| Intercept | 1.00 | 78659 | 76645 |
| Tooth Type (C) | 1.00 | 148529 | 82878 |
| Tooth Type (PM) | 1.00 | 157298 | 79091 |
| Isomere (U) | 1.00 | 191107 | 74358 |
| Position in Field (P) | 1.00 | 201647 | 74806 |
| Side (R) | 1.00 | 211635 | 77314 |
| DeW 2 | 1.00 | 171846 | 81439 |
| DeW 3 | 1.00 | 146969 | 84737 |
| DeW 4 | 1.00 | 148225 | 80433 |
| Sex (M) | 1.00 | 85792 | 75770 |
| Age (MA) | 1.00 | 104805 | 81459 |

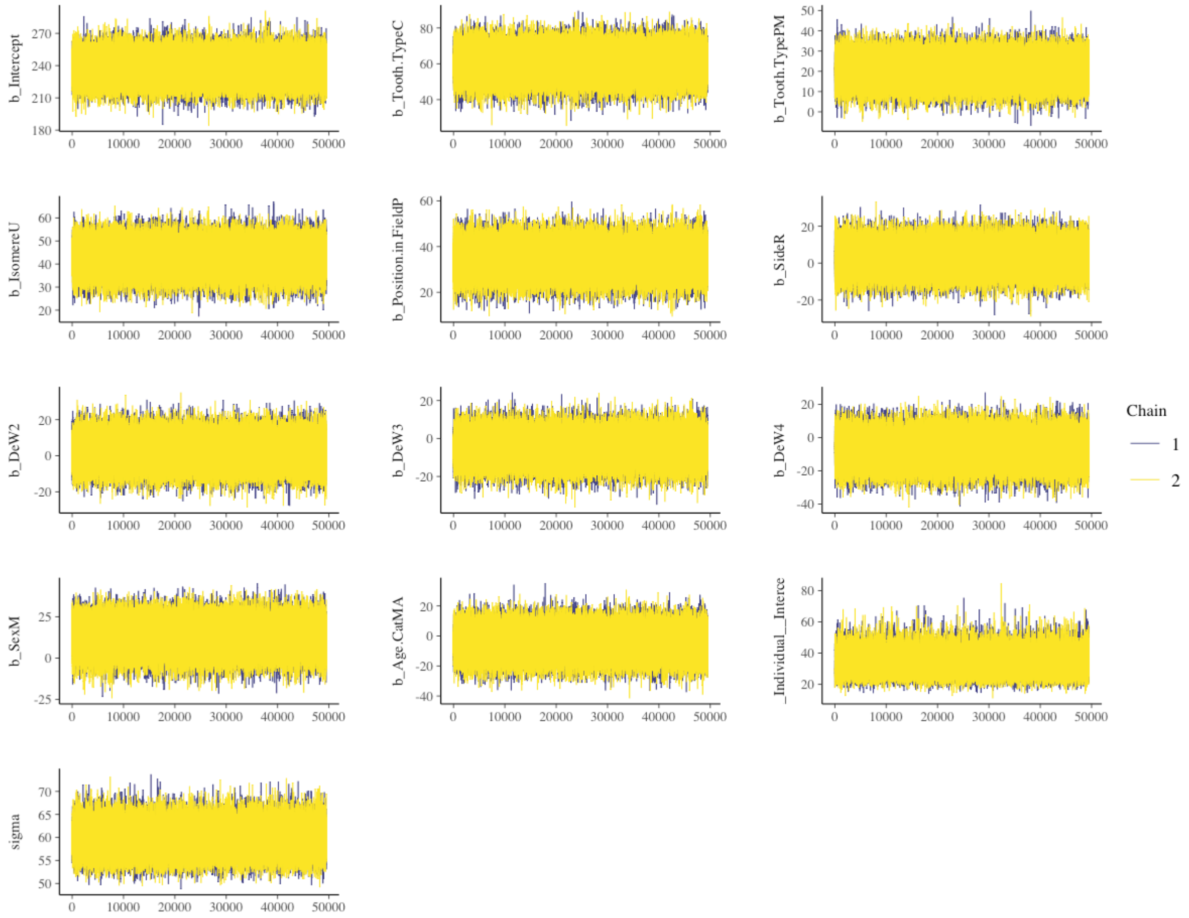

*Figure 4 Dentin volume M7 model Markov Chain Monte Carlo Trace for each parameter.*

#### 4.5 Pulp Volume

*Table 5 MCMC diagnostics for the Pulp Volume Model*

| Parameter | Rhat | Bulk ESS | Tail ESS |
|---|---|---|---|
| Random | | | |
| Individual | 1.00 | 34635 | 51369 |
| Tooth | 1.00 | 115536 | 74219 |
| Fixed | | | |
| Intercept | 1.00 | 49082 | 61916 |
| Tooth Type (C) | 1.00 | 124342 | 80927 |
| Tooth Type (PM) | 1.00 | 92785 | 80029 |
| Isomere (U) | 1.00 | 150137 | 72721 |
| Position in Field (P) | 1.00 | 141690 | 79633 |
| Side (R) | 1.00 | 163006 | 76389 |
| DeW 2 | 1.00 | 69292 | 75441 |
| DeW 3 | 1.00 | 58692 | 69693 |
| DeW 4 | 1.00 | 58662 | 70036 |
| Sex (M) | 1.00 | 37183 | 55066 |
| Age (MA) | 1.00 | 40888 | 57519 |

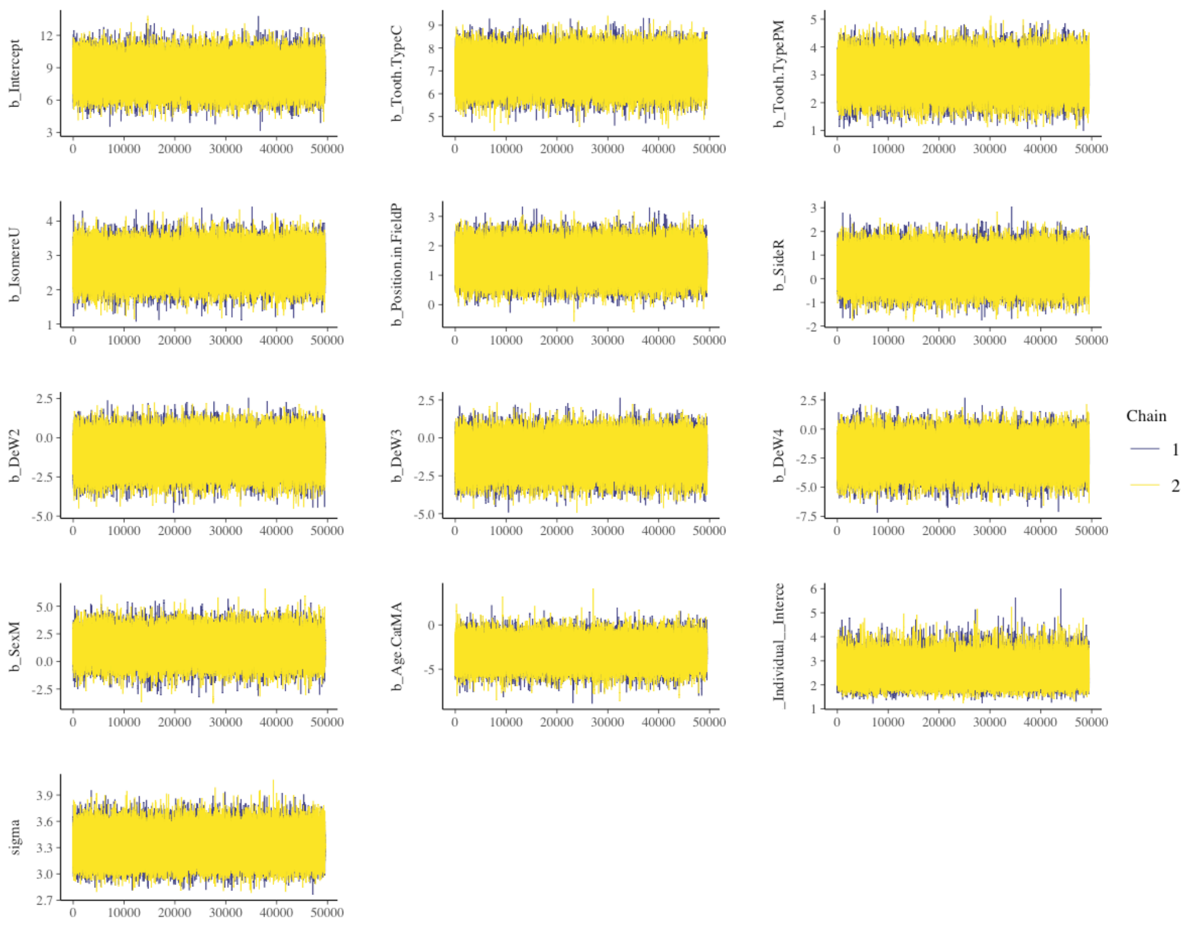

*Figure 5 Pulp volume M7 model Markov Chain Monte Carlo Trace for each parameter.*

#### 4.6 Crown Volume

*Table 6 MCMC diagnostics for the Crown Volume Model*

| Parameter | Rhat | Bulk ESS | Tail ESS |
|---|---|---|---|
| Random | | | |
| Individual | 1.00 | 37795 | 56333 |
| Tooth | 1.00 | 20901 | 78719 |
| Fixed | | | |
| Intercept | 1.00 | 79051 | 75635 |
| Tooth Type (C) | 1.00 | 159200 | 77716 |
| Tooth Type (PM) | 1.00 | 149815 | 80377 |
| Isomere (U) | 1.00 | 196510 | 72188 |
| Position in Field (P) | 1.00 | 164433 | 76228 |
| Side (R) | 1.00 | 193240 | 75935 |
| DeW 2 | 1.00 | 138491 | 84501 |
| DeW 3 | 1.00 | 116792 | 82796 |
| DeW 4 | 1.00 | 107532 | 76911 |
| Sex (M) | 1.00 | 79532 | 77574 |
| Age (MA) | 1.00 | 86565 | 78241 |

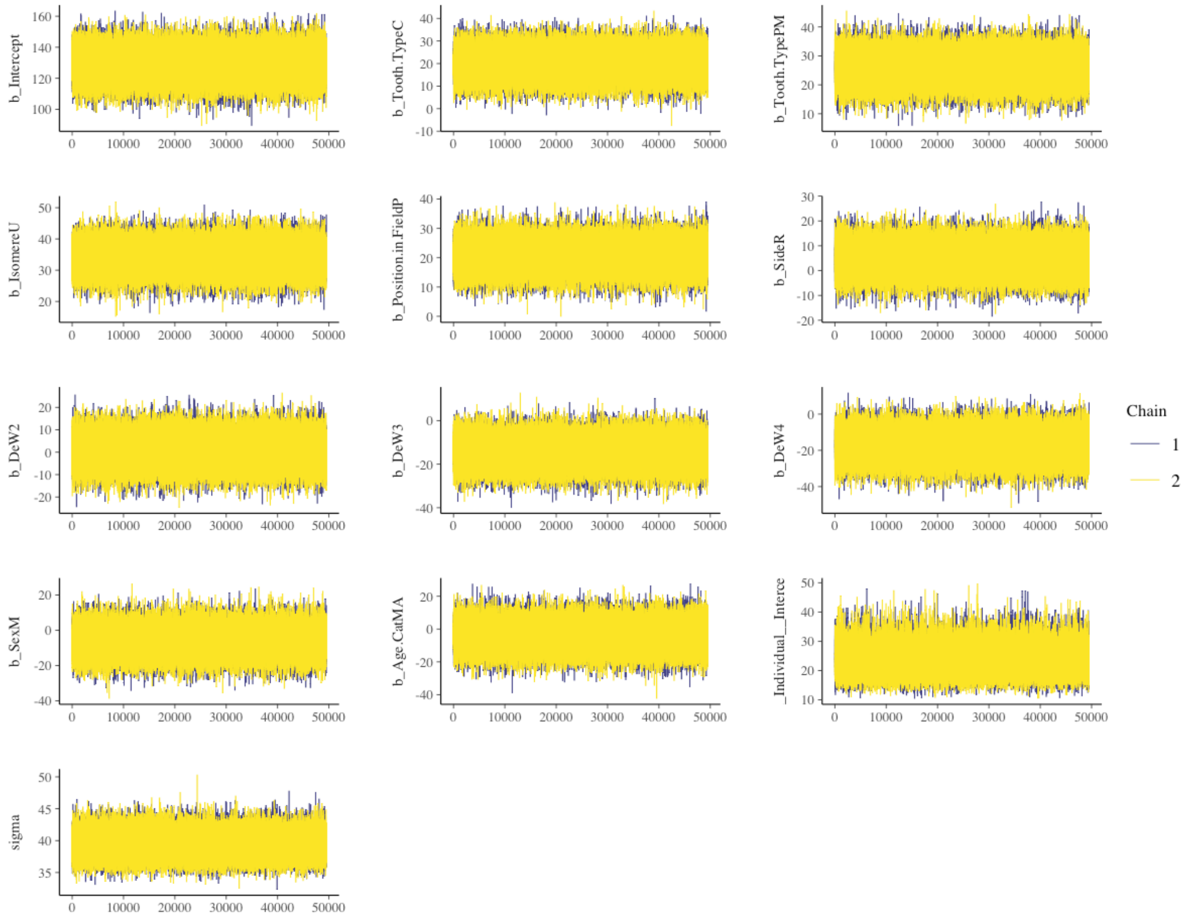

*Figure 6 Crown volume M7 model Markov Chain Monte Carlo Trace for each parameter.*

#### 4.7 Crown Surface Area

*Table 7 MCMC diagnostics for the Crown Surface Area Model*

| Parameter | Rhat | Bulk ESS | Tail ESS |
|---|---|---|---|
| Random | | | |
| Individual | 1.00 | 37610 | 58361 |
| Tooth | 1.00 | 132108 | 77743 |
| Fixed | | | |
| Intercept | 1.00 | 64444 | 72808 |
| Tooth Type (C) | 1.00 | 147572 | 80899 |
| Tooth Type (PM) | 1.00 | 125953 | 79352 |
| Isomere (U) | 1.00 | 188469 | 71957 |
| Position in Field (P) | 1.00 | 183305 | 79384 |
| Side (R) | 1.00 | 179016 | 72842 |
| DeW 2 | 1.00 | 104821 | 85440 |
| DeW 3 | 1.00 | 82555 | 80499 |
| DeW 4 | 1.00 | 79559 | 78245 |
| Sex (M) | 1.00 | 56270 | 68772 |
| Age (MA) | 1.00 | 59750 | 70403 |

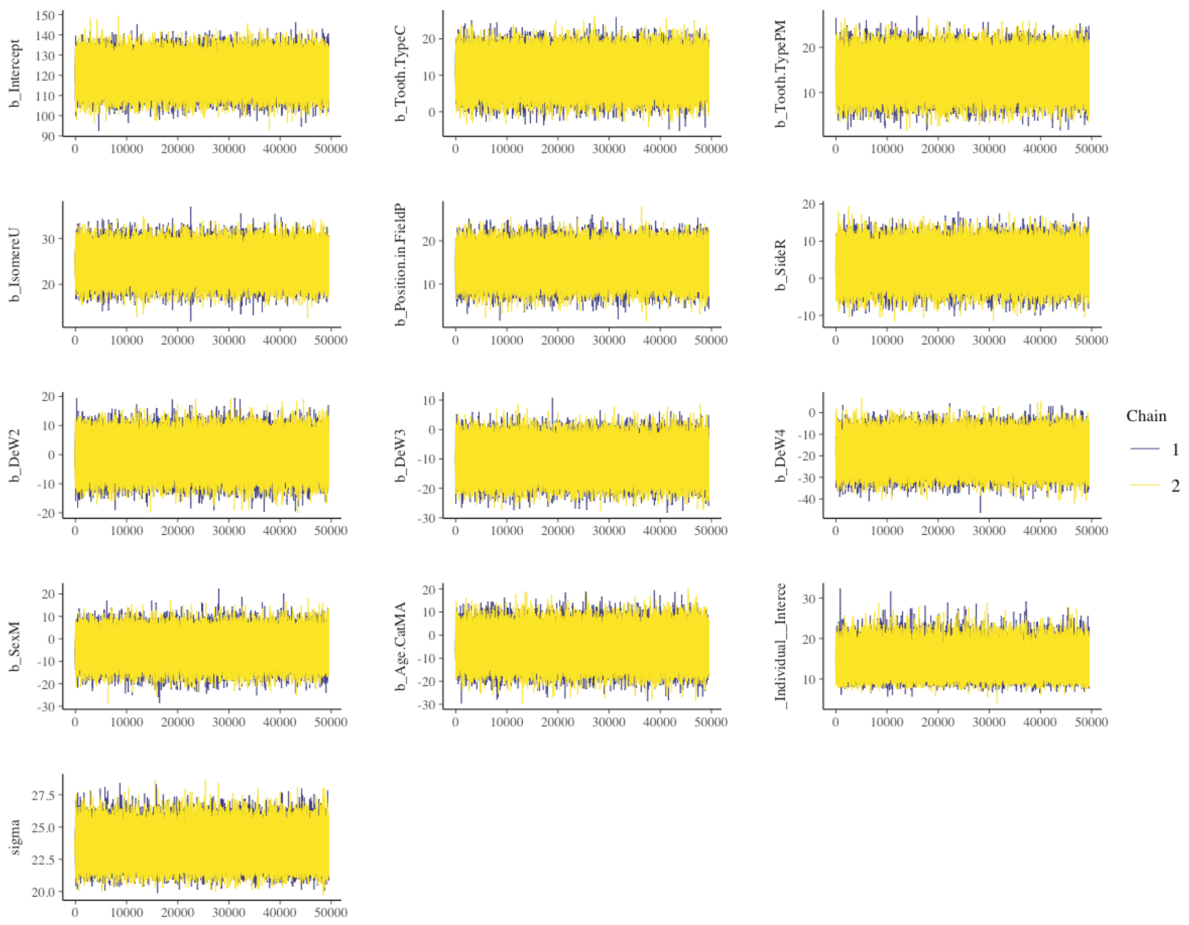

*Figure 7 Crown surface area M7 model Markov Chain Monte Carlo Trace for each parameter.*

#### 4.8 Root Volume

*Table 8 MCMC diagnostics for the Crown Volume Model*

| Parameter | Rhat | Bulk ESS | Tail ESS |
|---|---|---|---|
| Random | | | |
| Individual | 1.00 | 33140 | 47991 |
| Tooth | 1.00 | 94593 | 81497 |
| Fixed | | | |
| Intercept | 1.00 | 68953 | 71489 |
| Tooth Type (C) | 1.00 | 109615 | 78906 |
| Tooth Type (PM) | 1.00 | 132918 | 79258 |
| Isomere (U) | 1.00 | 143491 | 74215 |
| Position in Field (P) | 1.00 | 150251 | 76331 |
| Side (R) | 1.00 | 142145 | 74844 |
| DeW 2 | 1.00 | 127788 | 80750 |
| DeW 3 | 1.00 | 115742 | 78973 |
| DeW 4 | 1.00 | 117715 | 77057 |
| Sex (M) | 1.00 | 75665 | 75752 |
| Age (MA) | 1.00 | 80702 | 74836 |

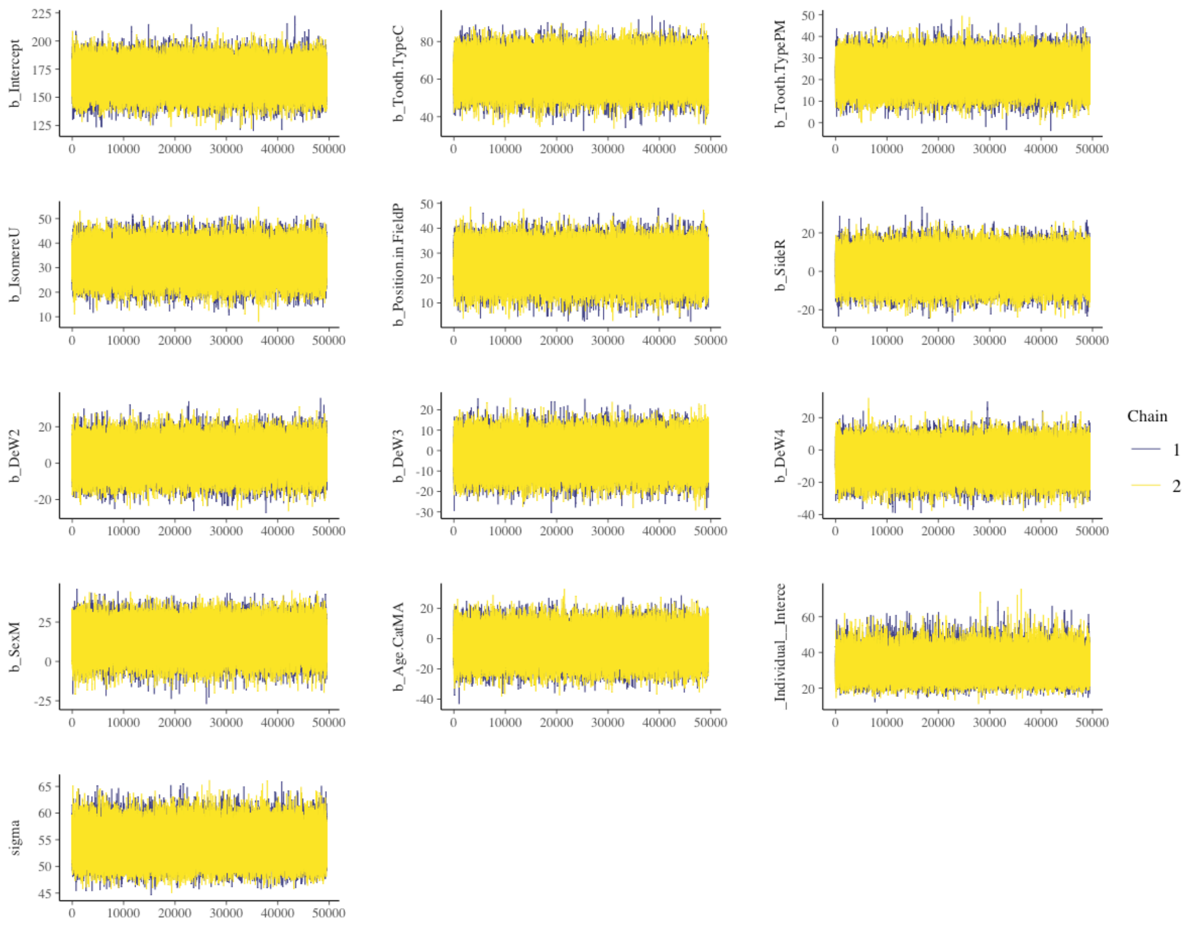

*Figure 8 Root volume M7 model Markov Chain Monte Carlo Trace for each parameter.*

#### 4.9 Root Surface Area

*Table 9 MCMC diagnostics for the Root Surface Area Model*

| Parameter | Rhat | Bulk ESS | Tail ESS |
|---|---|---|---|
| Random | | | |
| Individual | 1.00 | 35954 | 55484 |
| Tooth | 1.00 | 126119 | 74811 |
| Fixed | | | |
| Intercept | 1.00 | 64590 | 73325 |
| Tooth Type (C) | 1.00 | 157572 | 82661 |
| Tooth Type (PM) | 1.00 | 150311 | 82215 |
| Isomere (U) | 1.00 | 222785 | 69215 |
| Position in Field (P) | 1.00 | 191437 | 76705 |
| Side (R) | 1.00 | 185828 | 73444 |
| DeW 2 | 1.00 | 128966 | 84426 |
| DeW 3 | 1.00 | 106055 | 84453 |
| DeW 4 | 1.00 | 107267 | 84594 |
| Sex (M) | 1.00 | 55812 | 65971 |
| Age (MA) | 1.00 | 74814 | 77376 |

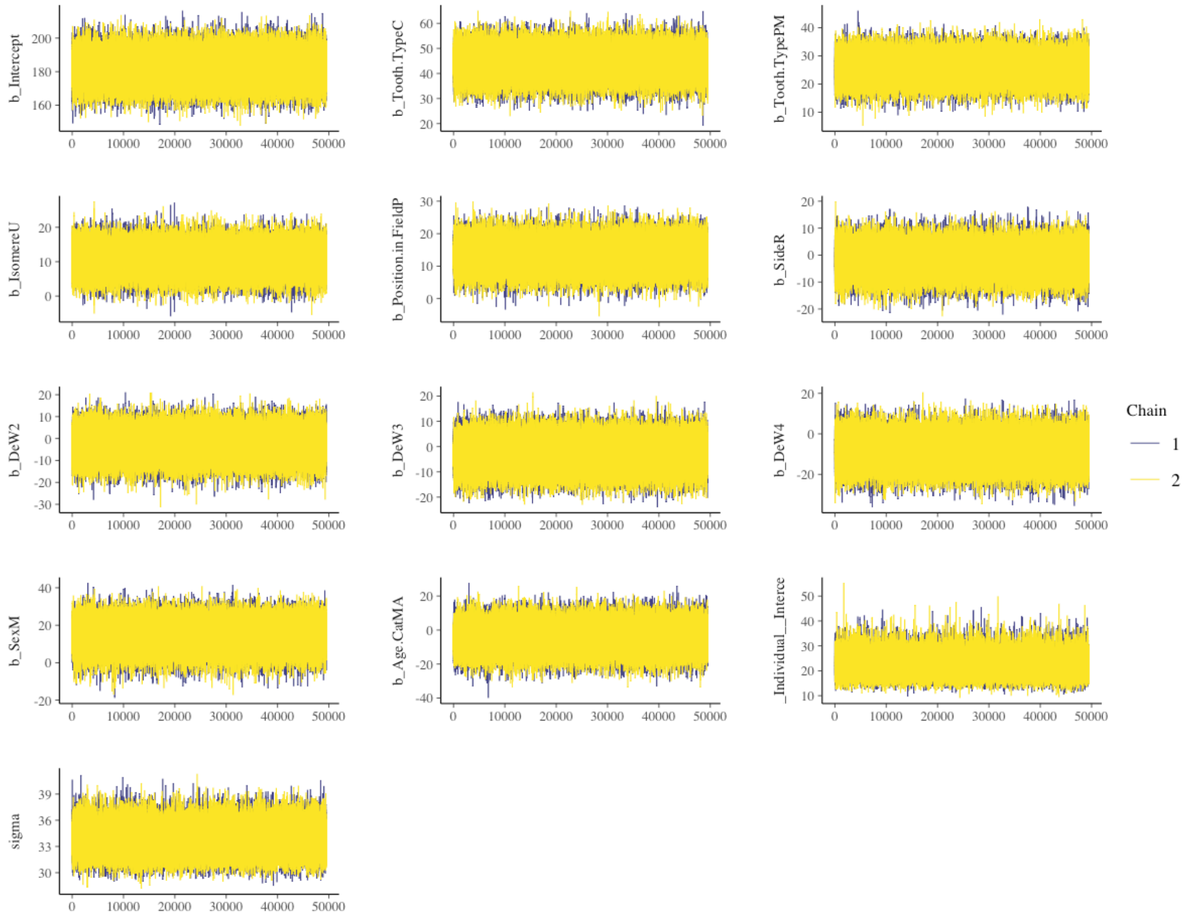

*Figure 9 Root surface area M7 model Markov Chain Monte Carlo Trace for each parameter.*

#### 4.10 Coronal Dentin Volume

*Table 10 MCMC diagnostics for the Coronal Dentin Volume Model*

| Parameter | Rhat | Bulk ESS | Tail ESS |
|---|---|---|---|
| Random | | | |
| Individual | 1.00 | 27377 | 25728 |
| Tooth | 1.00 | 112984 | 70612 |
| Fixed | | | |
| Intercept | 1.00 | 108878 | 82075 |
| Tooth Type (C) | 1.00 | 158141 | 78172 |
| Tooth Type (PM) | 1.00 | 154590 | 82736 |
| Isomere (U) | 1.00 | 218565 | 72219 |
| Position in Field (P) | 1.00 | 187651 | 78535 |
| Side (R) | 1.00 | 208482 | 75647 |
| DeW 2 | 1.00 | 114198 | 82338 |
| DeW 3 | 1.00 | 96490 | 81734 |
| DeW 4 | 1.00 | 96974 | 81864 |
| Sex (M) | 1.00 | 100516 | 76960 |
| Age (MA) | 1.00 | 111367 | 79993 |

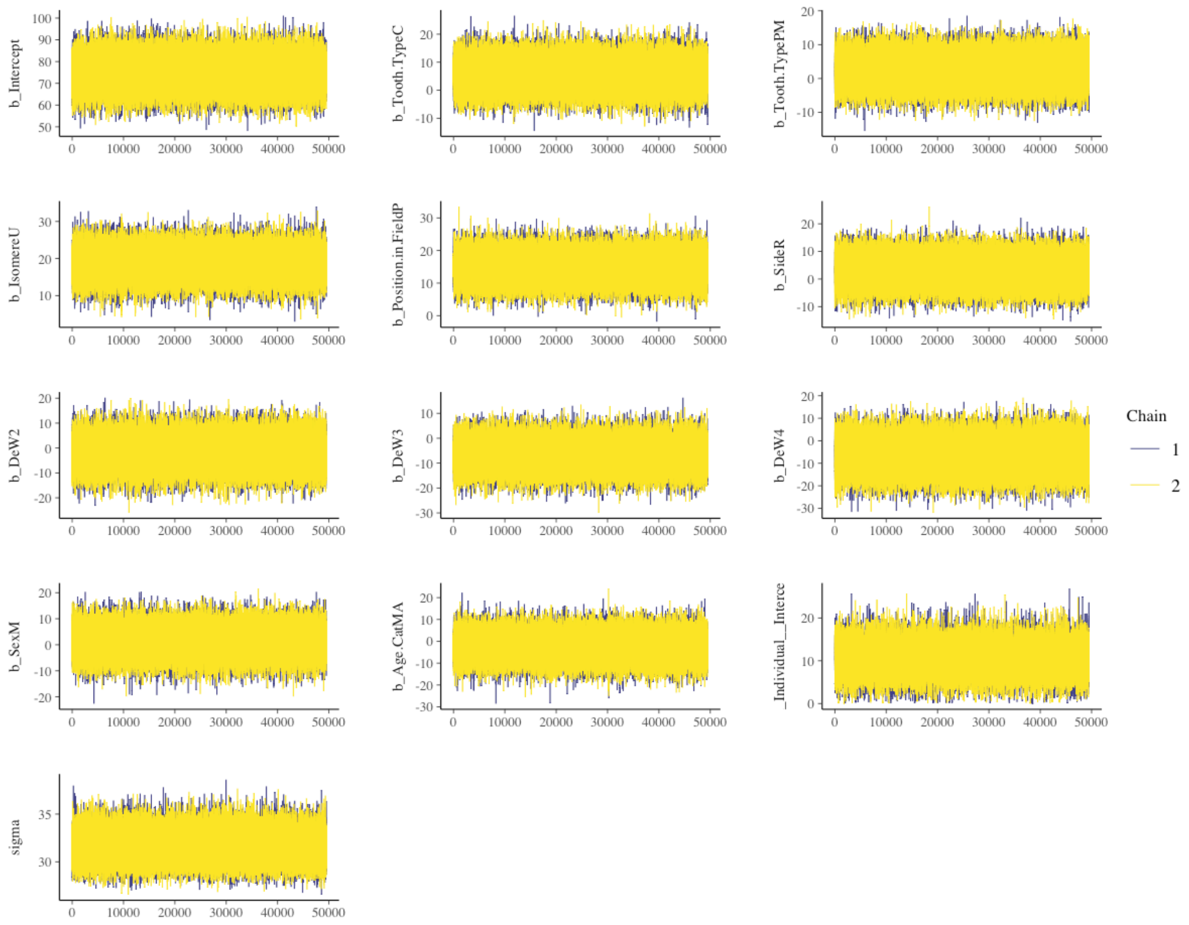

*Figure 10. Coronal dentin volume M7 model Markov Chain Monte Carlo Trace for each parameter.*

#### 4.11 Whole Tooth Volume

*Table 11 MCMC diagnostics for the Whole Tooth Volume Model*

| Parameter | Rhat | Bulk ESS | Tail ESS |
|---|---|---|---|
| Random | | | |
| Individual | 1.00 | 34248 | 52005 |
| Tooth | 1.00 | 83589 | 75338 |
| Fixed | | | |
| Intercept | 1.00 | 78692 | 73895 |
| Tooth Type (C) | 1.00 | 121392 | 82504 |
| Tooth Type (PM) | 1.00 | 130650 | 82452 |
| Isomere (U) | 1.00 | 127418 | 77306 |
| Position in Field (P) | 1.00 | 147163 | 76217 |
| Side (R) | 1.00 | 143877 | 74515 |
| DeW 2 | 1.00 | 135863 | 81067 |
| DeW 3 | 1.00 | 130447 | 80499 |
| DeW 4 | 1.00 | 134830 | 81378 |
| Sex (M) | 1.00 | 97352 | 74281 |
| Age (MA) | 1.00 | 107832 | 79117 |

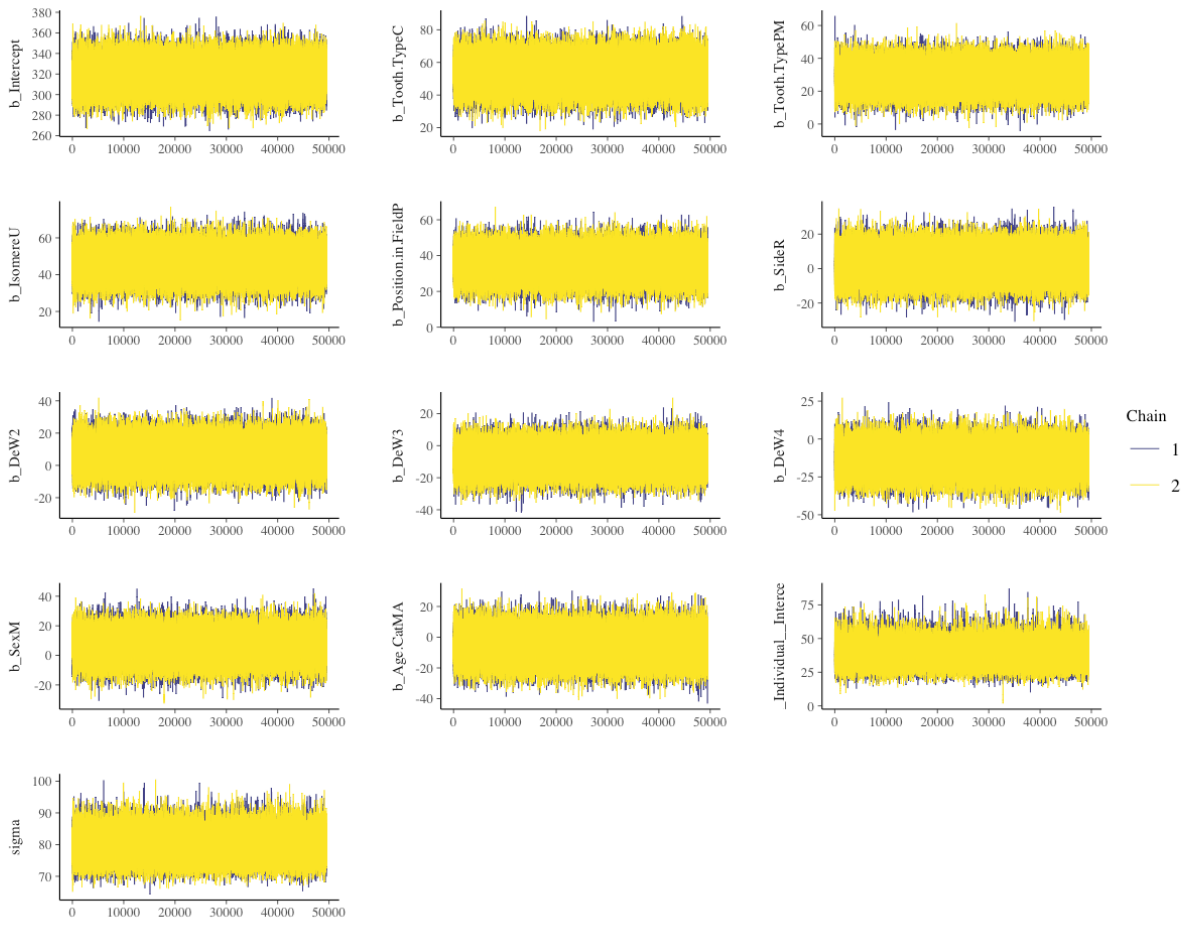

*Figure 11 Whole tooth volume M7 model Markov Chain Monte Carlo Trace for each parameter.*

#### 4.12 Whole Tooth Surface Area

*Table 12 MCMC diagnostics for the Whole Tooth Surface Area Model*

| Parameter | Rhat | Bulk ESS | Tail ESS |
|---|---|---|---|
| Random | | | |
| Individual | 1.00 | 32170 | 45919 |
| Tooth | 1.00 | 126815 | 83644 |
| Fixed | | | |
| Intercept | 1.00 | 52070 | 66284 |
| Tooth Type (C) | 1.00 | 145249 | 82477 |
| Tooth Type (PM) | 1.00 | 143536 | 81696 |
| Isomere (U) | 1.00 | 208780 | 71620 |
| Position in Field (P) | 1.00 | 185684 | 75784 |
| Side (R) | 1.00 | 188394 | 75409 |
| DeW 2 | 1.00 | 135963 | 81630 |
| DeW 3 | 1.00 | 111597 | 82299 |
| DeW 4 | 1.00 | 113162 | 78366 |
| Sex (M) | 1.00 | 63000 | 69481 |
| Age (MA) | 1.00 | 70627 | 73238 |

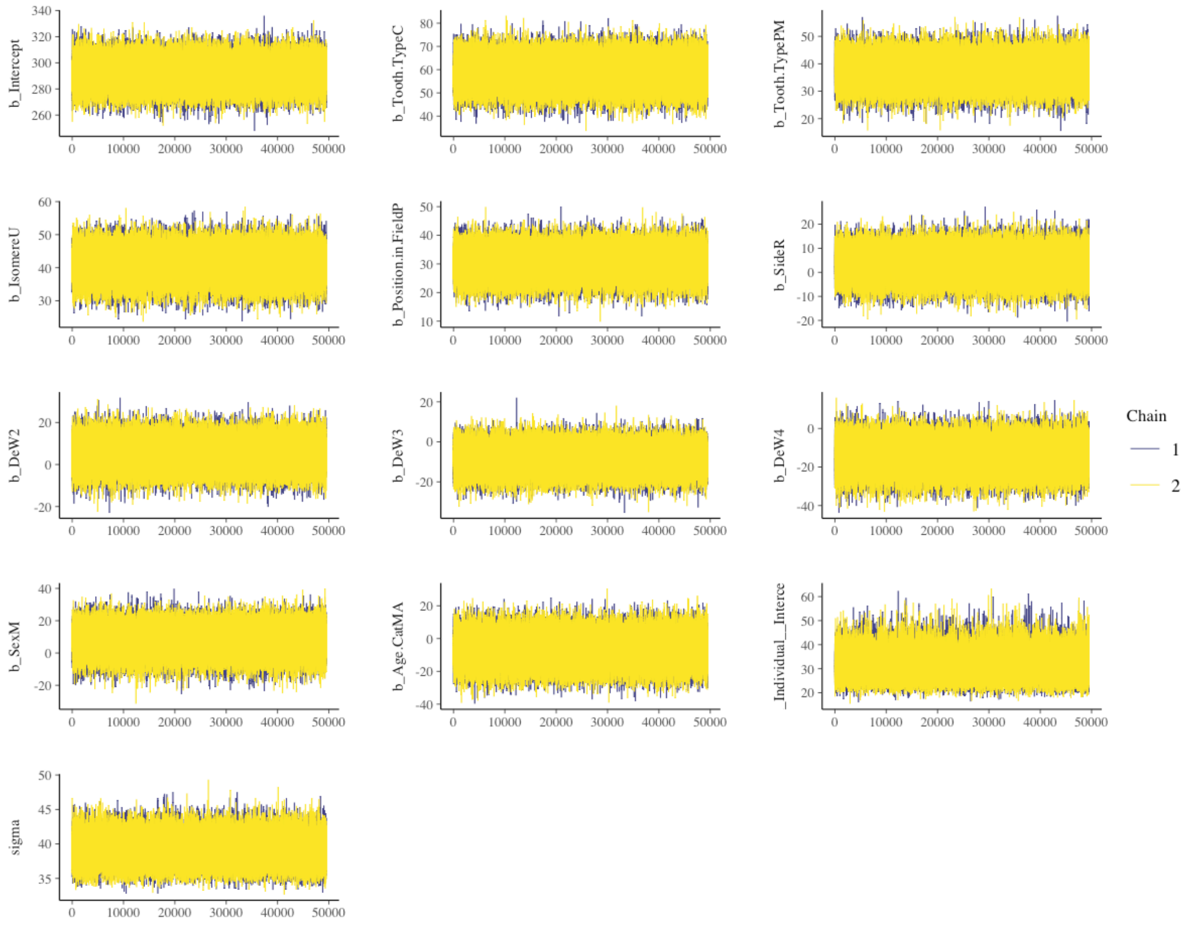

*Figure 12 Whole tooth surface area M7 model Markov Chain Monte Carlo Trace for each parameter.*

### 5 Supporting Information: Model Comparison

*Table 13 Leave-one-out (LOO) cross-validation results for each multilevel model*

| Model | $M_n$ | $M_1$ | $M_2$ | $M_3$ | $M_4$ | $M_5$ | $M_6$ | $M_7$ |
|---|---|---|---|---|---|---|---|---|
| ESA Loo | 3214.9 | 3175.8 | 3072.5 | 3037.4 | 3036.6 | 3024.6 | 3024.7 | 3024.5 |
| DSA Loo | 3400.4 | 3275.1 | 3198.0 | 3144.7 | 3145.9 | 3144.3 | 3143.5 | 3143.5 |
| EVol Loo | 3214.9 | 3175.8 | 3072.5 | 3037.4 | 3036.6 | 3024.6 | 3024.7 | 3024.5 |
| DVol Loo | 3529.8 | 3444.7 | 3374.5 | 3331.9 | 3332.9 | 3333.5 | 3332.8 | 3332.8 |
| Pvol Loo | 1814.5 | 1659.4 | 1615.8 | 1604.5 | 1606.0 | 1607.6 | 1608.0 | 1607.0 |
| CSALoo | 2919.4 | 2883.6 | 2809.4 | 2785.5 | 2786.2 | 2779.5 | 2779.7 | 2779.7 |
| RSA Loo | 3138.7 | 3015.1 | 3007.9 | 2999.7 | 3001.6 | 3002.7 | 3001.6 | 3002.3 |
| CVol Loo | 3226.3 | 3181.1 | 3113.0 | 3091.8 | 3092.3 | 3085.8 | 3085.7 | 3086.1 |
| RVol Loo | 3461.9 | 3348.0 | 3296.5 | 3275.8 | 3276.9 | 3278.2 | 3277.4 | 3277.4 |
| CDVol Loo | 2994.6 | 2990.5 | 2962.1 | 2946.8 | 2947.9 | 2949.4 | 2950.3 | 2951.0 |
| WTSA | 3361.2 | 3235.8 | 3138.2 | 3089.6 | 3090.3 | 3086.4 | 3086.2 | 3086.0 |
| WTVol Loo | 3671.5 | 3611.3 | 3548.6 | 3508.0 | 3508.7 | 3503.9 | 3503.6 | 3503.6 |

### 6 Supporting Information: Model Outputs

#### 6.1 Enamel Volume

*Table 14 Enamel Volume MLM results summary. Posterior means (mm) with the 95% credibility interval for each of the parameters in the model. Variance estimates are given for each of the random parameters (individual and tooth) as a standard deviation. This is followed by the leave-one-out (LOO) cross-validation estimate and the Variance Partition Coefficient (VPC) for level 2.*

| Parameter | $M_n$ | $M_1$ | $M_2$ | $M_3$ | $M_4$ | $M_5$ | $M_6$ | $M_7$ |
|---|---|---|---|---|---|---|---|---|
| Random | | | | | | | | |
| Individual | 34.28 (24.68, 46.87) | 34.72 (25.33, 47.10) | 35.72 (26.63, 47.79) | 35.98 (26.98, 48.12) | 35.90 (26.97, 47.98) | 30.39 (22.05, 41.37) | 29.92 (21.73, 40.82) | 35.32 (32.39, 38.58) |
| Tooth | 49.15 (45.24, 53.53) | 45.87 (42.12, 50.02) | 38.44 (35.29, 41.96 | 36.18 (33.19, 39.51) | 36.09 (33.12, 39.42) | 35.32 (32.41, 38.59) | 35.32 (32.41, 38.60) | 29.68 (21.50, 40.71) |
| Fixed | | | | | | | | |
| Intercept | 229.60 (215.97, 243.18) | 215.89 (201.47, 230.31) | 194.55 (179.73, 209.36) | 180.99 (165.68, 196.43) | 179.57 (163.97, 195.06) | 187.11 (170.41, 203.74) | 190.02 (172.02, 207.66) | 191.45 (173.03, 209.56) |
| Tooth Type (C) | | 26.56 (5.87, 14.98) | 30.19 (19.97, 40.24) | 22.37 (12.14, 32.49) | 22.41 (12.23, 32.48) | 20.37 (10.36, 30.36) | 20.36 (10.23, 30.41) | 20.39 (10.31, 30.47) |
| Tooth Type (PM) | | 20.86 (10.93, 30.70) | 23.36 (14.61, 31.97) | 24.92 (16.64, 33.07) | 25.29 (17.04, 33.45) | 20.79 (12.29, 29.17) | 20.82 (12.35, 29.23) | 20.87 (12.37, 29.27) |
| Isomere (U) | | | 39.48 (31.39, 47.49) | 40.23 (32.58, 47.76) | 40.22 (32.56, 47.82) | 41.45 (33.83, 48.95) | 41.45 (33.85, 48.88) | 41.44 (33.85, 48.89) |
| Position in Field (P) | | | | 23.58 (15.39, 31.74) | 23.70 (15.53, 31.79) | 23.72 (15.64, 31.74) | 23.72 (15.68, 31.68) | 23.72 (15.64, 31.74) |
| Side (R) | | | | | 6.29 | 7.10 | 7.28 | 7.27 |

| | | | | | | | | |
|---|---|---|---|---|---|---|---|---|
| | | | | | (-3.80, 16.38) | (-2.80, 16.94) | (-2.59, 17.15) | (-2.64, 17.16) |
| DeW (2) | | | | | | 8.05 (-3.66, 19.78) | 8.00 (-3.77, 19.71) | 8.00 (-3.60, 19.73) |
| DeW (3) | | | | | | -6.57 (-17.45, 4.39) | -6.45 (-17.36, 4.52) | -6.36 (-17.28, 4.64) |
| DeW (4) | | | | | | -23.73 (-37.54, -9.77) | -23.51 (-37.30, -9.68) | 23.31 (-37.13, -9.44) |
| Sex (M) | | | | | | | -6.66 (-21.58, 8.44) | -6.61 (-21.32, 8.44) |
| Age (MA) | | | | | | | | -5.79 (-21.39, 10.03) |
| Loo | 3214.9 | 3175.8 | 3072.5 | 3037.4 | 3036.6 | 3024.6 | 3024.7 | 3024.5 |
| ICC | 0.327 | 0.364 | 0.463 | 0.497 | 0.497 | 0.425 | 0.418 | 0.414 |

#### 6.2 Enamel Surface Area

*Table 15 Enamel Surface Area MLM results summary. Posterior means (mm) with the 95% credibility interval for each of the parameters in the model. Variance estimates are given for each of the random parameters (individual and tooth) as a standard deviation. This is followed by the leave-one-out (LOO) cross-validation estimate and the Variance Partition Coefficient (VPC) for level 2.*

| Parameter | $M_n$ | $M_1$ | $M_2$ | $M_3$ | $M_4$ | $M_5$ | $M_6$ | $M_7$ |
|---|---|---|---|---|---|---|---|---|
| Random | | | | | | | | |
| Individual | 34.28 (24.68, 46.87) | 34.72 (25.33, 47.10) | 35.72 (26.63, 47.79) | 35.98 (26.98, 48.12) | 35.90 (26.97, 47.98) | 30.39 (22.05, 41.37) | 29.92 (21.73, 40.82) | 35.32 (32.39, 38.58) |
| Tooth | 49.15 (45.24, 53.53) | 45.87 (42.12, 50.02) | 38.44 (35.29, 41.96 | 36.18 (33.19, 39.51) | 36.09 (33.12, 39.42) | 35.32 (32.41, 38.59) | 35.32 (32.41, 38.60) | 29.68 (21.50, 40.71) |
| Fixed | | | | | | | | |
| Intercept | 229.60 (215.97, 243.18) | 215.89 (201.47, 230.31) | 194.55 (179.73, 209.36) | 180.99 (165.68, 196.43) | 179.57 (163.97, 195.06) | 187.11 (170.41, 203.74) | 190.02 (172.02, 207.66) | 191.45 (173.03, 209.56) |
| Tooth Type (C) | | 26.56 (5.87, 14.98) | 30.19 (19.97, 40.24) | 22.37 (12.14, 32.49) | 22.41 (12.23, 32.48) | 20.37 (10.36, 30.36) | 20.36 (10.23, 30.41) | 20.39 (10.31, 30.47) |
| Tooth Type (PM) | | 20.86 (10.93, 30.70) | 23.36 (14.61, 31.97) | 24.92 (16.64, 33.07) | 25.29 (17.04, 33.45) | 20.79 (12.29, 29.17) | 20.82 (12.35, 29.23) | 20.87 (12.37, 29.27) |
| Isomere (U) | | | 39.48 (31.39, 47.49) | 40.23 (32.58, 47.76) | 40.22 (32.56, 47.82) | 41.45 (33.83, 48.95) | 41.45 (33.85, 48.88) | 41.44 (33.85, 48.89) |
| Position in Field (P) | | | | 23.58 (15.39, 31.74) | 23.70 (15.53, 31.79) | 23.72 (15.64, 31.74) | 23.72 (15.68, 31.68) | 23.72 (15.64, 31.74) |
| Side (R) | | | | | 6.29 (-3.80, 16.38) | 7.10 (-2.80, 16.94) | 7.28 (-2.59, 17.15) | 7.27 (-2.64, 17.16) |
| DeW (2) | | | | | | 8.05 | 8.00 | 8.00 |

| | | | | | | | | |
|---|---|---|---|---|---|---|---|---|
| | | | | | | (-3.66, 19.78) | (-3.77, 19.71) | (-3.60, 19.73) |
| DeW (3) | | | | | | -6.57 (-17.45, 4.39) | -6.45 (-17.36, 4.52) | -6.36 (-17.28, 4.64) |
| DeW (4) | | | | | | -23.73 (-37.54, -9.77) | -23.51 (-37.30, -9.68) | 23.31 (-37.13, -9.44) |
| Sex (M) | | | | | | | -6.66 (-21.58, 8.44) | -6.61 (-21.32, 8.44) |
| Age (MA) | | | | | | | | -5.79 (-21.39, 10.03) |
| Loo | 3214.9 | 3175.8 | 3072.5 | 3037.4 | 3036.6 | 3024.6 | 3024.7 | 3024.5 |
| ICC | 0.327 | 0.364 | 0.463 | 0.497 | 0.497 | 0.425 | 0.418 | 0.414 |

#### 6.3 Dentin Volume

*Table 16 Dentin Volume MLM results summary. Posterior means (mm) with the 95% credibility interval for each of the parameters in the model. Variance estimates are given for each of the random parameters (individual and tooth) as a standard deviation. This is followed by the leave-one-out (LOO) cross-validation estimate and the Variance Partition Coefficient (VPC) for level 2.*

| Parameter | $M_n$ | $M_1$ | $M_2$ | $M_3$ | $M_4$ | $M_5$ | $M_6$ | $M_7$ |
|---|---|---|---|---|---|---|---|---|
| Random | | | | | | | | |
| Individual | 30.32 (15.65, 46.63) | 33.74 (21.58, 48.80) | 35.62 (24.29, 50.09) | 36.50 (25.53, 50.79) | 36.39 (25.47, 50.46) | 36.78 (25.82, 51.05) | 33.84 (22.87, 47.88) | 33.62 (22.56, 47.76) |
| Tooth | 84.18 (77.36, 91.75) | 72.48 (66.13, 79.56) | 64.11 (58.23, 70.61) | 59.52 (54.17, 65.51) | 59.56 (54.20, 65.61) | 59.49 (54.11, 65.53) | 59.49 (54.07, 65.50) | 59.49 (54.08, 65.54) |
| Fixed | | | | | | | | |
| Intercept | 301.9 (287.33, 316.79) | 284.59 (268.22, 300.94) | 261.44 (243.93, 278.84) | 239.24 (220.67, 258.04) | 238.72 (219.90, 257.52) | 242.38 (221.07, 263.87) | 236.66 (215.07, 258.94) | 237.69 (215.56, 260.13) |
| Tooth Type | | 56.87 (41.14, 72.29) | 65.01 (49.67, 79.93) | 60.67 (45.95, 75.09) | 60.64 (45.82, 75.22) | 60.30 (45.52, 74.61) | 60.28 (45.40, 74.75) | 60.29 (45.56, 74.69) |
| Tooth Type 2 | | 14.93 (1.61, 28.04) | 18.17 (5.63, 30.62 | 21.95 (9.94, 33.82 | 22.08 (9.85, 33.97) | 20.86 (8.64, 32.79 | 20.78 (8.57, 32.74) | 20.84 (8.63, 32.71) |
| Isomere | | | 40.48 (28.37, 52.32) | 42.54 (30.98, 53.74) | 42.50 (30.94, 53.86) | 42.75 (31.25, 53.99) | 42.72 (31.21, 54.00) | 42.72 (31.15, 53.97) |
| Position in Field | | | | 34.21 (22.42, 45.87) | 34.19 (22.42, 45.87) | 34.35 (22.49, 46.15) | 34.37 (22.48, 46.24) | 34.40 (22.51, 46.13) |
| Side | | | | | 2.96 (-10.61, 16.59) | 3.07 (-10.74, 16.71) | 2.85 (-10.89, 16.57) | 2.82 (-10.88 16.53) |
| DeW 2 | | | | | | 2.39 | 2.69 | 2.61 |

| | | | | | | | | |
|---|---|---|---|---|---|---|---|---|
| | | | | | | (-12.27, 16.89) | (-11.80, 17.26) | (-11.95, 17.17) |
| DeW 3 | | | | | | -4.73 (-18.20, 8.85) | -5.02 (-18.39, 8.42) | -4.93 (-18.38, 8.55) |
| DeW 4 | | | | | | -7.02 (-22.80, 8.77) | -7.49 (-23.27, 8.27) | -7.23 (-22.94, 8.56) |
| Sex | | | | | | | 12.79 (-4.16, 29.09) | 12.86 (-3.99, 29.13) |
| Age | | | | | | | | -4.64 (-21.29, 12.10) |
| Loo | 3529.8 | 3444.7 | 3374.5 | 3331.9 | 3332.9 | 3333.5 | 3332.8 | 3332.8 |
| ICC | 0.115 | 0.178 | 0.236 | 0.273 | 0.272 | 0.277 | 0.244 | 0.242 |

#### 6.4 Dentin Surface Area

*Table 17 Dentin Surface Area MLM results summary. Posterior means (mm) with the 95% credibility interval for each of the parameters in the model. Variance estimates are given for each of the random parameters (individual and tooth) as a standard deviation. This is followed by the leave-one-out (LOO) cross-validation estimate and the Variance Partition Coefficient (VPC) for level 2.*

| Parameter | $M_n$ | $M_1$ | $M_2$ | $M_3$ | $M_4$ | $M_5$ | $M_6$ | $M_7$ |
|---|---|---|---|---|---|---|---|---|
| Random | | | | | | | | |
| Individual | 34.47 (23.04, 48.83 | 36.89 (26.42, 50.51) | 37.91 (27.86, 51.35) | 38.46 (28.47, 51.81) | 38.39 (28.44, 51.65 | 38.18 (28.34, 51.26) | 36.87 (27.07, 50.09) | 35.90 (26.00, 49.20) |
| Tooth | 67.32 (61.93, 73.32) | 54.20 (49.47, 59.46) | 47.39 (43.17, 52.09) | 43.20 (39.41, 47.44) | 43.27 (39.47, 47.52) | 43.01 (39.24, 47.23) | 42.99 (39.23, 47.23) | 42.99 (39.22 47.23) |
| Fixed | | | | | | | | |
| Intercept | 372.44 (357.73, 387.13) | 349.88 (334.17, 365.60) | 328.90 (312.10, 345.40) | 308.08 (290.94, 325.30) | 307.88 (290.73, 325.13) | 311.48 (291.74, 331.18) | 307.27 (286.77, 328.26) | 309.06 (288.31, 330.08) |
| Tooth Type (C) | | 63.15 (49.41, 76.54) | 70.61 (57.77, 83.01) | 64.05 (51.86, 75.88) | 63.98 (51.76, 75.82) | 63.05 (50.92, 74.86) | 63.03 (50.92, 74.91) | 63.07 (50.91, 74.94) |
| Tooth Type (PM) | | 24.84 (13.50, 35.94) | 28.55 (18.14, 38.70 | 32.40 (22.65, 41.97 | 32.41 (22.60, 42.06) | 30.20 (20.20, 40.03) | 30.11 (20.16, 39.96) | 30.16 (20.13, 39.89) |
| Isomere (U) | | | 36.06 (26.36, 45.52) | 37.53 (28.5, 46.37) | 37.49 (28.56, 46.31) | 37.94 (28.93, 46.80) | 37.95 (29.03, 46.75) | 37.95 (29.04, 46.71) |
| Position in Field (P) | | | | 33.15 (23.78, 42.52) | 33.18 (23.80, 42.50) | 33.44 (24.02, 42.82) | 33.43 (24.10, 42.77) | 33.45 (23.99, 42.84) |
| Side (R) | | | | | 1.13 (-10.28, 12.57) | 1.40 (-9.92, 12.72) | 1.22 (-10.17, 12.54) | 1.27 (-10.07, 12.57) |
| DeW 2 | | | | | | 7.25 | 7.41 | 7.37 |

| | | | | | | | | |
|---|---|---|---|---|---|---|---|---|
| | | | | | | (-5.53, 20.08) | (-5.41, 20.23) | (-5.45, 20.18) |
| DeW 3 | | | | | | -4.58 (-16.43, 7.32) | -4.78 (-16.62, 7.10) | -4.71 (-16.62, 7.22) |
| DeW 4 | | | | | | -11.10 (-25.61, 3.42) | -11.49 (-25.93, 2.93) | -11.21 (-25.59, 3.30) |
| Sex (M) | | | | | | | 9.50 (-7.14, 25.63) | 9.90 (-6.67, 26.07) |
| Age (MA) | | | | | | | | -7.81 (-24.42, 9.10) |
| Loo | 3400.4 | 3275.1 | 3198.0 | 3144.7 | 3145.9 | 3144.3 | 3143.5 | 3143.5 |
| ICC | 0.208 | 0.317 | 0.390 | 0.442 | 0.441 | 0.441 | 0.424 | 0.411 |

#### 6.5 Pulp Volume

*Table 18 Pulp Volume MLM results summary. Posterior means (mm) with the 95% credibility interval for each of the parameters in the model. Variance estimates are given for each of the random parameters (individual and tooth) as a standard deviation. This is followed by the leave-one-out (LOO) cross-validation estimate and the Variance Partition Coefficient (VPC) for level 2.*

| Parameter | $M_n$ | $M_1$ | $M_2$ | $M_3$ | $M_4$ | $M_5$ | $M_6$ | $M_7$ |
|---|---|---|---|---|---|---|---|---|
| Random | | | | | | | | |
| Individual | 2.72 (1.88, 3.80) | 2.90 (2.13, 3.93) | 2.95 (2.18, 3.97) | 2.95 (2.19, 3.98) | 2.95 (2.19, 3.97) | 2.90 (2.14, 3.91) | 2.87 (2.11, 3.90) | 2.54 (1.82, 3.50) |
| Tooth | 4.77 (4.39, 5.19) | 3.65 (3.36, 3.97) | 3.38 (3.11, 3.68) | 3.31 (3.04, 3.61) | 3.32 (3.05 , 3.61) | 3.31 (3.04, 3.61) | 3.31 (3.04, 3.61) | 3.31 (3.04, 3.61) |
| Fixed | | | | | | | | |
| Intercept | 11.89 (10.77, 13.01) | 8.94 (7.69, 10.17) | 7.63 (6.35, 8.91) | 6.89 (5.54, 8.22) | 6.80 (5.46, 8.16) | 8.29 (6.14, 10.44) | 7.83 (5.51, 10.14) | 8.43 (6.19, 10.65) |
| Tooth Type (C) | | 7.90 (6.77, 9.04) | 7.90 (6.85, 8.96) | 7.16 (6.05, 8.26) | 7.16 (6.06, 8.28) | 7.01 (5.89, 8.13) | 6.99 (5.86, 8.11) | 7.02 (5.89, 8.14) |
| Tooth Type (PM) | | 3.43 (2.51, 4.36) | 3.44 (2.58, 4.29) | 3.44 (2.60, 4.27) | 3.46 (2.62, 4.31) | 2.95 (1.97, 3.92) | 2.90 (1.91, 3.88) | 2.98 (2.01, 3.96) |
| Isomere (U) | | | 2.61 (1.84, 3.37) | 2.61 (1.86, 3.36) | 2.61 (1.85, 3.35) | 2.73 (1.97, 3.49) | 2.74 (1.98, 3.50) | 2.72 (1.96, 3.49) |
| Position in Field (P) | | | | 1.49 (0.65, 2.33) | 1.50 (0.66, 2.34) | 1.45 (0.61, 2.29) | 1.45 (0.61, 2.29) | 1.45 (0.61, 2.29) |
| Side (R) | | | | | 0.41 (-0.66, 1.47) | 0.47 (-0.60, 1.55) | 0.44 (-0.64, 1.52) | 0.43 (-0.64, 1.50) |
| DeW (2) | | | | | | -1.09 (-2.79, 0.62) | -1.13 (-2.85, 0.60) | -1.04 (-2.75, 0.68) |
| DeW (3) | | | | | | -1.23 (-2.96, 0.50) | -1.34 (-3.11, 0.40) | -1.18 (-2.92, 0.56) |
| DeW (4) | | | | | | -2.45 (-4.66, | -2.63 (-4.89, -0.40) | -2.27 (-4.49, -0.02) |

| | | | | | | | | |
|---|---|---|---|---|---|---|---|---|
| | | | | | | -0.24) | | |
| Sex (M) | | | | | | | 1.28 (-1.00, 3.53) | 1.33 (-0.70, 3.36) |
| Age (MA) | | | | | | | | -3.07 (-5.36, -0.77) |
| Loo | 1814.5 | 1659.4 | 1615.8 | 1604.5 | 1606.0 | 1607.6 | 1608.0 | 1607.0 |
| ICC | 0.245 | 0.388 | 0.432 | 0.442 | 0.442 | 0.434 | 0.429 | 0.370 |

#### 6.6 Crown Volume

*Table 19 Crown volume MLM results summary. Posterior means (mm) with the 95% credibility interval for each of the parameters in the model. Variance estimates are given for each of the random parameters (individual and tooth) as a standard deviation. This is followed by the leave-one-out (LOO) cross-validation estimate and the Variance Partition Coefficient (VPC) for level 2.*

| Parameter | $M_n$ | $M_1$ | $M_2$ | $M_3$ | $M_4$ | $M_5$ | $M_6$ | $M_7$ |
|---|---|---|---|---|---|---|---|---|
| Random | | | | | | | | |
| Individual | 25.23 (16.69, 35.87) | 26.08 (17.91, 36.44) | 27.01 (19.19, 37.34) | 27.28 (19.56, 37.41) | 27.36 (19.62, 37.51) | 23.76 (16.21, 33.40) | 23.42 (15.95, 33.03) | 23.41 (15.92, 33.12) |
| Tooth | 50.42 (46.36, 54.90) | 46.53 (42.72, 50.78) | 41.33 (37.92, 45.15) | 39.80 (36.50, 43.48) | 39.78 (36.47, 43.42) | 39.30 (36.02, 42.94) | 39.30 (36.03, 42.92) | 39.30 (36.00, 42.95) |
| Fixed | | | | | | | | |
| Intercept | 158.55 (147.70, 169.39) | 143.17 (131.18, 155.17) | 125.38 (112.73, 138.13) | 114.01 (100.59, 127.57) | 113.14 (99.57, 126.85) | 125.14 (109.37, 140.96) | 127.87 (110.89, 144.66) | 128.89 (111.64, 145.99) |
| Tooth Type (C) | | 25.17 (13.55. 36.68) | 27.68 (16.81, 38.34) | 21.22 (10.24, 32.01) | 21.21 (10.19, 32.08) | 20.05 (9.38, 30.76) | 20.07 (9.26, 30.79) | 20.09 (9.30, 30.77) |
| Tooth Type (PM) | | 25.97 (15.94, 35.82) | 27.97 (18.70, 37.11) | 29.28 (20.36, 38.08) | 29.51 (20.45, 38.35) | 25.73 (16.56, 34.81) | 25.81 (16.51, 34.91) | 25.88 (16.67, 35.02) |
| Isomere (U) | | | 32.97 (24.47, 41.35) | 33.43 (25.13, 41.56) | 33.44 (25.13, 41.60) | 34.12 (25.84, 42.32) | 34.11 (25.8, 42.22) | 34.11 (25.86, 42.25) |
| Position in Field (P) | | | | 19.90 (11.07, 28.61) | 19.95 (11.08, 28.80) | 20.55 (11.79, 29.24) | 20.56 (11.75, 29.24) | 20.57 (11.7, 29.33) |
| Side (R) | | | | | 4.07 (-6.72, 14.75) | 4.44 (-6.20, 15.05) | 4.66 (-5.96, 15.31) | 4.69 (-5.94, 15.30) |
| DeW (2) | | | | | | 0.84 (-26.16, | 0.74 | 0.68 |

| | | | | | | | | |
|---|---|---|---|---|---|---|---|---|
| | | | | | | -3.27) | (-11.60, 13.06) | (-11.64, 12.96) |
| DeW (3) | | | | | | -14.74 (-26.16, -3.27) | -14.52 (-25.90, -3.09) | -14.42 (-25.81, -3.01) |
| DeW (4) | | | | | | -18.35 (-32.51, -4.07) | -17.90 (-32.03, -3.67) | -17.58 (-31.79, -3.40) |
| Sex (M) | | | | | | | -6.42 (-20.09, 7.56) | -6.41 (-20.11, 7.52) |
| Age (MA) | | | | | | | | -4.23 (-18.69, 10.38) |
| Loo | 3226.3 | 3181.1 | 3113.0 | 3091.8 | 3092.3 | 3085.8 | 3085.7 | 3086.1 |
| ICC | 0.200 | 0.239 | 0.299 | 0.320 | 0.321 | 0.268 | 0.262 | 0.262 |

#### 6.7 Crown Surface Area

*Table 20 Crown surface area MLM results summary. Posterior means (mm) with the 95% credibility interval for each of the parameters in the model. Variance estimates are given for each of the random parameters (individual and tooth) as a standard deviation. This is followed by the leave-one-out (LOO) cross-validation estimate and the Variance Partition Coefficient (VPC) for level 2.*

| Parameter | $M_n$ | $M_1$ | $M_2$ | $M_3$ | $M_4$ | $M_5$ | $M_6$ | $M_7$ |
|---|---|---|---|---|---|---|---|---|
| Random | | | | | | | | |
| Individual | 16.66 (11.43, 23.38) | 17.00 (11.90, 23.59) | 17.58 (12.70, 24.02) | 17.76 (12.90, 24.21) | 17.79 (12.98, 24.21) | 14.21 (9.66, 20.07) | 14.24 (9.68, 20.02) | 14.24 (9.63, 20.14) |
| Tooth | 30.10 (27.69, 32.77) | 28.23 (25.94, 30.78) | 24.81 (22.80, 27.06) | 23.79 (8.50, 19.95) | 23.78 (21.85, 25.93) | 23.50 (21.59, 25.64) | 23.51 (21.59, 25.62) | 23.50 (21.59, 25.64) |
| Fixed | | | | | | | | |
| Intercept | 138.10 (131.11, 145.01) | 127.78 (120.04, 135.59) | 115.59 (107.43, 123.76) | 108.05 (99.47, 116.71) | 107.37 (98.67, 116.05) | 118.25 (106.91, 129.33) | 119.96 (108.01, 131.64) | 120.92 (108.86, 132.84) |
| Tooth Type (C) | | 17.42 (9.38, 25.40) | 18.30 (3.63, 11.17) | 12.30 (4.94, 19.61) | 18.37 (4.99, 19.63) | 10.50 (3.17, 17.79) | 10.55 (3.22, 17.89) | 10.59 (3.27, 17.93) |
| Tooth Type (PM) | | 17.05 (10.38, 23.69) | 17.72 (11.79, 23.64) | 18.18 (12.42, 23.90) | 18.37 (12.59, 24.10) | 13.93 (7.79, 20.06) | 14.07 (7.96, 20.22) | 14.20 (8.08, 20.33) |
| Isomere (U) | | | 23.54 (18.11, 28.98) | 23.68 (18.41, 28.91) | 23.67 (18.43, 28.85) | 24.54 (19.33, 29.74) | 24.52 (19.30, 29.69) | 24.48 (19.31, 29.62) |
| Position in Field (P) | | | | 14.22 (8.50, 19.95) | 14.28 (8.58, 19.94) | 14.47 (8.83, 20.10) | 14.47 (8.76, 20.17) | 14.47 (8.81, 20.16) |
| Side (R) | | | | | 3.05 (-4.11, 10.20) | 3.45 (-3.66, 10.53) | 3.63 (-3.38, 10.64) | 3.66 (-3.40, 10.71) |
| DeW (2) | | | | | | -0.14 (-9.34, 9.09) | -0.17 (-9.38, 9.13) | -0.13 (-9.34, 9.08) |

| | | | | | | | | |
|---|---|---|---|---|---|---|---|---|
| DeW (3) | | | | | | -10.91 (-19.67, -2.05) | -10.61 (-19.47, -1.80) | -10.34 (-19.14, -1.480 |
| DeW (4) | | | | | | -19.73 (-31.01, -8.29) | -19.14 (-30.47, -7.77) | -18.52 (-29.83, -7.16) |
| Sex (M) | | | | | | | -4.40 (-14.46, 5.65) | -4.29 (-14.36, 5.86) |
| Age (MA) | | | | | | | | -4.90 (-15.81, 6.10) |
| Loo | 2919.4 | 2883.6 | 2809.4 | 2785.5 | 2786.2 | 2779.5 | 2779.7 | 2779.7 |
| ICC | 0.234 | 0.266 | 0.334 | 0.358 | 0.359 | 0.268 | 0.268 | 0.269 |

#### 6.8 Root Volume

*Table 21 Root volume MLM results summary. Posterior means (mm) with the 95% credibility interval for each of the parameters in the model. Variance estimates are given for each of the random parameters (individual and tooth) as a standard deviation. This is followed by the leave-one-out (LOO) cross-validation estimate and the Variance Partition Coefficient (VPC) for level 2.*

| Parameter | $M_n$ | $M_1$ | $M_2$ | $M_3$ | $M_4$ | $M_5$ | $M_6$ | $M_7$ |
|---|---|---|---|---|---|---|---|---|
| Random | | | | | | | | |
| Individual | 30.19 (17.77, 44.99) | 33.43 (22.87, 46.88) | 34.57 (24.36, 47.92) | 34.88 (24.71, 48.21) | 34.79 (24.67, 47.93) | 35.48 (25.14, 48.85) | 32.45 (22.08, 45.82) | 32.22 (21.84, 45.53) |
| Tooth | 74.77 (68.77, 81.42) | 61.29 (55.84, 67.34) | 56.04 (50.98, 61.71) | 54.02 (49.24, 59.36) | 54.07 (49.27, 59.45) | 54.05 (49.25, 59.44) | 54.03 (49.22, 59.40) | 54.04 (49.28, 59.39) |
| Fixed | | | | | | | | |
| Intercept | 223.55 (209.53, 237.60) | 203.00 (187.75, 218.32) | 185.09 (168.69, 201.46) | 169.85 (152.55, 187.25) | 169.59 (152.01, 187.22) | 171.77 (151.34, 192.28) | 165.76 (144.87, 186.88) | 166.77 (145.60, 188.37) |
| Tooth Type (C) | | 63.72 (48.84, 78.21) | 69.38 (55.04, 83.23) | 64.31 (50.18, 78.02) | 64.21 (50.07, 77.91) | 63.76 (49.57, 77.64) | 63.76 (49.61, 77.54) | 63.79 (49.75, 77.44) |
| Tooth Type (PM) | | 19.44 (7.24, 31.46) | 21.93 (10.31, 33.35) | 24.17 (12.89, 35.32) | 24.19 (12.76, 35.40) | 23.06 (11.51, 34.51) | 22.99 (11.46, 34.33) | 23.01 (11.48, 34.36) |
| Isomere (U) | | | 31.47 (20.57, 42.19) | 32.16 (21.56, 42.62) | 32.14 (21.63, 42.55) | 32.38 (21.71, 42.84) | 32.39 (21.77, 42.87) | 32.40 (21.81, 42.79) |
| Position in Field (P) | | | | 25.02 (14.04, 35.98) | 25.06 (14.17, 35.88) | 25.02 (14.03, 35.95) | 25.07 (13.94, 36.06) | 25.06 (13.97, 36.16) |
| Side (R) | | | | | 1.73 (-11.20, 14.70) | 1.85 (-11.22, 14.91) | 1.51 (-11.49, 14.47) | 1.54 (-11.42, 14.48) |
| DeW (2) | | | | | | 2.76 | 3.00 | 2.92 |

| | | | | | | | | |
|---|---|---|---|---|---|---|---|---|
| | | | | | | (-11.30, 16.91) | (-11.13, 17.13) | (-11.12, 17.03) |
| DeW (3) | | | | | | -2.00 (-15.05, 11.02) | -2.37 (-15.32, 10.63) | -2.32 (-15.24, 10.67) |
| DeW (4) | | | | | | -7.01 (-22.55, 8.51) | -7.33 (-22.88, 8.19) | -7.11 (-22.57, 8.36) |
| Sex (M) | | | | | | | 13.59 (-3.17, 29.69) | 13.72 (-3.01, 29.80) |
| Age (MA) | | | | | | | | -4.52 (-20.83, 11.88) |
| Loo | 3461.9 | 3348.0 | 3296.5 | 3275.8 | 3276.9 | 3278.2 | 3277.4 | 3277.4 |
| ICC | 0.140 | 0.229 | 0.276 | 0.294 | 0.293 | 0.301 | 0.265 | 0.262 |

#### 6.9 Root Surface Area

*Table 22 Root surface area MLM results summary. Posterior means (mm) with the 95% credibility interval for each of the parameters in the model. Variance estimates are given for each of the random parameters (individual and tooth) as a standard deviation. This is followed by the leave-one-out (LOO) cross-validation estimate and the Variance Partition Coefficient (VPC) for level 2.*

| Parameter | $M_n$ | $M_1$ | $M_2$ | $M_3$ | $M_4$ | $M_5$ | $M_6$ | $M_7$ |
|---|---|---|---|---|---|---|---|---|
| Random | | | | | | | | |
| Individual | 22.74 (15.33, 32.19) | 24.28 (17.46, 33.21) | 24.36 (17.57, 33.28) | 24.48 (17.71, 33.41) | 24.50 (17.74, 33.45) | 25.11 (18.20, 34.24) | 22.28 (15.55, 31.05) | 22.15 (15.36, 31.09) |
| Tooth | 43.40 (39.91, 47.30) | 34.90 (32.02, 38.10) | 34.37 (31.49, 37.55) | 33.82 (31.01, 36.96) | 33.86 (31.03, 37.03) | 33.81 (30.97, 36.95) | 33.81 (31.01, 36.95) | 33.81 (30.99, 36.95) |
| Fixed | | | | | | | | |
| Intercept | 215.87 (206.39, 225.44 | 195.37 (184.81, 205.84) | 190.13 (179.07, 201.27) | 182.67 (170.88, 194.61) | 183.00 (171.07, 195.03) | 186.79 (171.77, 201.93) | 180.66 (165.19, 196.52) | 181.79 (166.01, 197.75) |
| Tooth Type (C) | | 49.22 (39.45, 58.74) | 49.62 (39.98, 59.04) | 44.71 (34.75, 54.41) | 44.69 (34.78, 54.44) | 44.37 (34.32, 54.21) | 44.30 (34.33, 54.12) | 44.33 (34.32, 54.16) |
| Tooth Type (PM) | | 26.44 (18.33, 34.48) | 26.67 (18.67, 34.51) | 27.34 (19.44, 35.09) | 27.21 (19.32, 35.00) | 25.89 (17.70, 33.99) | 25.72 (17.44, 33.87) | 25.77 (17.52, 33.95) |
| Isomere (U) | | | 10.22 (2.99, 17.42) | 10.25 (3.11, 17.42) | 10.26 (3.06, 17.42) | 10.67 (3.45, 17.82) | 10.69 (3.55, 17.85) | 10.68 (3.50, 17.83) |
| Position in Field (P) | | | | 13.63 (5.87, 21.30) | 13.59 (5.84, 21.32) | 13.30 (5.51, 21.09) | 13.31 (5.59, 21.12) | 13.33 (5.58, 21.12) |
| Side (R) | | | | | -1.69 (-11.18, 7.80) | -1.50 (-11.02, 8.00) | -1.93 (-11.42 7.55) | -1.93 (-11.44, 7.64) |
| DeW (2) | | | | | | -3.00 | -2.89 | -2.96 |

| | | | | | | | | |
|---|---|---|---|---|---|---|---|---|
| | | | | | | (-14.33, 8.38) | (-14.26, 8.55) | (-14.30, 8.45) |
| DeW (3) | | | | | | -2.39 (-12.95, 8.24) | -3.05 (-13.69, 7.63) | -2.95 (-13.57, 7.74) |
| DeW (4) | | | | | | -8.25 (-21.44, 4.99) | -8.89 (6.69, -22.05) | -8.53 (-21.72, 4.75) |
| Sex (M) | | | | | | | 14.41 (0.42, 27.68) | 14.53 (0.35, 27.75) |
| Age (MA) | | | | | | | | -4.83 (-18.78, 9.35) |
| Loo | 3138.7 | 3015.1 | 3007.9 | 2999.7 | 3001.6 | 3002.7 | 3001.6 | 3002.3 |
| ICC | 0.215 | 0.326 | 0.334 | 0.344 | 0.344 | 0.356 | 0.303 | 0.300 |

#### 6.10 Coronal Dentin Volume

*Table 23 Coronal dentin volume MLM results summary. Posterior means (mm) with the 95% credibility interval for each of the parameters in the model. Variance estimates are given for each of the random parameters (individual and tooth) as a standard deviation. This is followed by the leave-one-out (LOO) cross-validation estimate and the Variance Partition Coefficient (VPC) for level 2.*

| Parameter | $M_n$ | $M_1$ | $M_2$ | $M_3$ | $M_4$ | $M_5$ | $M_6$ | $M_7$ |
|---|---|---|---|---|---|---|---|---|
| Random | | | | | | | | |
| Individual | 8.70 (1.44, 15.39) | 8.95 (1.83, 15.48) | 9.80 (3.36, 16.05) | 10.21 (4.26, 16.24) | 10.24 (4.28, 16.30) | 9.77 (3.47, 15.99) | 9.98 (3.58, 16.25) | 10.21 (3.80, 16.52) |
| Tooth | 34.69 (31.88, 37.77) | 34.30 (31.52, 37.36) | 32.51 (29.85, 35.47) | 31.57 (28.98, 34.45) | 31.57 (29.01, 34.42) | 31.59 (29.00, 34.48) | 31.59 (29.03, 34.46) | 31.59 (29.00, 34.47) |
| Fixed | | | | | | | | |
| Intercept | 91.01 (85.84, 96.17) | 87.36 (80.77, 93.97) | 78.05 (70.55, 85.52) | 70.18 (61.90, 78.50) | 69.59 (61.16, 78.06) | 75.10 (63.07, 87.12) | 74.65 (62.04, 87.16) | 75.01 (62.29, 87.74) |
| Tooth Type (C) | | 11.70 (2.39, 20.92) | 12.00 (3.03, 20.85) | 6.27 (-2.98, 15.44) | 6.29 (-2.90, 15.38) | 5.84 (-3.29, 14.99) | 5.81 (-3.40, 15.06) | 5.86 (-3.41, 15.05) |
| Tooth Type (PM) | | 3.27 (-4.53, 11.03) | 3.42 (-4.07, 10.90) | 3.89 (-3.44, 11.21) | 4.07 (-3.27, 11.40) | 2.49 (-5.20, 10.20) | 2.43 (-5.27, 10.06) | 2.54 (-5.21, 10.24) |
| Isomere (U) | | | 18.35 (11.35, 25.28) | 18.49 (11.72, 25.25) | 18.47 (11.74, 25.17) | 18.69 (11.92, 25.43) | 18.70 (11.93, 25.45) | 18.68 (11.96, 25.42) |
| Position in Field (P) | | | | 14.60 (7.26, 21.90) | 14.64 (7.36, 21.94) | 14.89 (7.54, 22.25) | 14.88 (7.52, 22.26) | 14.88 (7.49, 22.20) |
| Side (R) | | | | | 3.00 (-5.82, 11.86) | 3.23 (-5.61, 12.08) | 3.09 (-5.73, 11.90) | 3.12 (-5.80, 12.07) |
| DeW (2) | | | | | | -1.60 | -1.58 | -1.61 |

| | | | | | | | | |
|---|---|---|---|---|---|---|---|---|
| | | | | | | (-12.34, 9.18) | (-12.36. 9.15) | (-12.29, 9.19) |
| DeW (3) | | | | | | -6.65 (-16.48, 3.24) | -6.79 (-16.74, 3.18) | -6.63 (-16.66, 3.40) |
| DeW (4) | | | | | | -6.72 (-18.71, 5.38) | -6.93 (-19.17, 5.31) | -6.45 (-18.79, 6.07) |
| Sex (M) | | | | | | | 1.34 (-8.13, 10.68) | 1.30 (-8.18, 10.68) |
| Age (MA) | | | | | | | | -2.13 (-12.34, 8.13) |
| Loo | 2994.6 | 2990.5 | 2962.1 | 2946.8 | 2947.9 | 2949.4 | 2950.3 | 2951.0 |
| ICC | 0.059 | 0.064 | 0.083 | 0.095 | 0.095 | 0.087 | 0.091 | 0.095 |

#### 6.11 Whole Tooth Volume

*Table 24 Whole tooth volume MLM results summary. Posterior means (mm) with the 95% credibility interval for each of the parameters in the model. Variance estimates are given for each of the random parameters (individual and tooth) as a standard deviation. This is followed by the leave-one-out (LOO) cross-validation estimate and the Variance Partition Coefficient (VPC) for level 2.*

| Parameter | $M_n$ | $M_1$ | $M_2$ | $M_3$ | $M_4$ | $M_5$ | $M_6$ | $M_7$ |
|---|---|---|---|---|---|---|---|---|
| Random | | | | | | | | |
| Individual | 32.00 (11.07, 52.51) | 36.25 (19.66, 55.08) | 39.06 (24.71, 56.63) | 40.51 (26.76, 57.67) | 40.47 (26.75, 57.47) | 39.35 (25.76, 56.26) | 38.68 (24.98, 55.63) | 38.11 (24.21, 55.27) |
| Tooth | 107.08 (98.44, 116.73) | 96.27 (87.89, 105.64) | 86.23 (78.22, 95.17) | 80.29 (72.67, 88.83) | 80.33 (72.74, 88.80) | 79.59 (72.03, 88.01) | 79.56 (72.08, 87.95) | 79.60 (72.04, 88.05) |
| Fixed | | | | | | | | |
| Intercept | 382.23 (365.00, 399.37) | 364.15 (345.33, 383.05) | 339.52 (318.95, 360.00) | 315.58 (293.44, 338.10) | 315.08 (292.72, 337.62) | 320.27 (295.93, 344.78) | 317.58 (292.38, 342.96) | 318.86 (293.31, 344.37) |
| Tooth Type (C) | | 47.26 (29.86, 64.44) | 55.09 (37.82, 72.00) | 53.43 (36.69, 69.76) | 53.38 (36.58, 69.71) | 53.49 (36.63, 69.94) | 53.50 (36.81, 69.99) | 53.49 (36.65, 69.91) |
| Tooth Type (PM) | | 21.68 (6.52, 36.81) | 25.66 (10.90, 40.28) | 30.31 (15.82, 44.57) | 30.35 (15.97, 44.54) | 28.95 (14.42, 43.13) | 28.93 (14.53, 43.06) | 28.92 (14.40, 43.26) |
| Isomere (U) | | | 42.84 (28.17, 57.20) | 45.86 (31.67, 59.65) | 45.80 (31.54, 59.64) | 46.47 (32.24, 60.27) | 46.48 (32.45, 60.18) | 46.47 (32.46, 60.30) |
| Position in Field (P) | | | | 34.98 (20.86, 48.87) | 35.00 (20.97, 48.86) | 35.46 (21.39, 49.29) | 35.47 (21.44, 49.32) | 35.47 (21.44, 49.31) |
| Side (R) | | | | | 2.70 (-12.79, 18.15) | 2.99 (-12.40, 18.32) | 2.79 (-12.83, 18.34) | 2.87 (-12.57, 18.31) |
| DeW (2) | | | | | | 6.78 | 6.96 | 6.92 |

| | | | | | | | | |
|---|---|---|---|---|---|---|---|---|
| | | | | | | (-9.27, 22.90) | (-9.08, 23.01) | (-9.18, 22.95) |
| DeW (3) | | | | | | -7.92 (-22.82, 7.04) | -8.10 (-22.92, 6.79) | -8.08 (-22.97, 6.84) |
| DeW (4) | | | | | | -13.13 (-29.83, 3.68) | -13.41 (-30.27, 3.58) | -13.23 (-30.10, 3.54) |
| Sex (M) | | | | | | | 6.07 (-11.06, 23.15) | 6.21 (-10.90, 23.09) |
| Age (MA) | | | | | | | | -5.71 (-22.94, 11.82) |
| Loo | 3671.5 | 3611.3 | 3548.6 | 3508.0 | 3508.7 | 3503.9 | 3503.6 | 3503.6 |
| ICC | 0.082 | 0.124 | 0.170 | 0.203 | 0.202 | 0.196 | 0.191 | 0.186 |

#### 6.12 Whole Tooth Surface Area

*Table 25 Whole tooth surface area MLM results summary. Posterior means (mm) with the 95% credibility interval for each of the parameters in the model. Variance estimates are given for each of the random parameters (individual and tooth) as a standard deviation. This is followed by the leave-one-out (LOO) cross-validation estimate and the Variance Partition Coefficient (VPC) for level 2.*

| Parameter | $M_n$ | $M_1$ | $M_2$ | $M_3$ | $M_4$ | $M_5$ | $M_6$ | $M_7$ |
|---|---|---|---|---|---|---|---|---|
| Random | | | | | | | | |
| Individual | 29.14 (18.63, 42.11) | 31.76 (22.35, 44.06) | 32.97 (24.04, 44.81) | 33.47 (24.67, 45.15) | 33.47 (24.70, 45.09) | 32.53 (23.86, 44.13) | 31.82 (23.21, 43.23) | 31.22 (22.61, 42.57) |
| Tooth | 63.21 (58.13, 68.84) | 50.86 (46.45, 55.78) | 42.90 (39.10, 47.16) | 39.42 (36.01, 43.26) | 39.43 (36.00, 43.27) | 39.07 (35.65, 42.89) | 39.08 (35.67, 42.90) | 39.08 (35.68, 42.91) |
| Fixed | | | | | | | | |
| Intercept | 353.45 (340.60, 366.30) | 328.86 (314.74, 343.08) | 305.71 (291.09, 320.38) | 287.09 (271.96, 302.43) | 286.57 (271.09, 301.96) | 293.67 (276.07, 311.43) | 290.48 (271.72, 309.40) | 292.08 (273.05, 311.01) |
| Tooth Type (C) | | 58.69 (45.58, 71.39) | 67.02 (55.06, 78.45) | 60.00 (48.45, 71.09) | 59.95 (48.43, 71.15) | 58.95 (47.59, 70.02) | 58.89 (47.59, 69.92) | 58.92 (47.55, 70.02) |
| Tooth Type (PM) | | 32.07 (21.15, 42.84) | 36.93 (27.18, 46.47) | 40.32 (31.17, 49.21) | 40.46 (31.30, 49.41) | 37.51 (28.14 46.68) | 37.42 (28.06, 46.59) | 37.44 (28.15, 46.63) |
| Isomere (U) | | | 39.10 (30.11, 47.91) | 40.33 (32.04, 48.47) | 40.31 (32.06, 48.43) | 40.91 (32.61, 49.11) | 40.94 (32.72, 49.05) | 40.92 (32.59, 49.07) |
| Position in Field (P) | | | | 29.97 (21.20, 38.62) | 30.02 (21.21, 38.79) | 30.39 (21.66, 39.04) | 30.41 (21.67, 39.11) | 30.42 (21.68, 39.11) |
| Side (R) | | | | | 3.05 (-7.57, 13.70) | 3.43 (-7.2, 14.10) | 3.26 (-7.34, 13.86) | 3.31 (-7.45, 14.01) |
| DeW (2) | | | | | | 4.92 | 5.02 | 4.97 |

| | | | | | | | | |
|---|---|---|---|---|---|---|---|---|
| | | | | | | (-7.35, 17.14) | (-7.17, 17.24) | (-7.33, 17.23) |
| DeW (3) | | | | | | -8.68 (-20.09, 2.83) | -8.91 (-20.24, 2.52) | -8.84 (-20.27, 2.62) |
| DeW (4) | | | | | | -14.55 (-28.53, -0.57) | -15.00 (-28.94, -0.92) | -14.72 (-28.88, -0.71) |
| Sex (M) | | | | | | | 7.30 (-8.33, 22.65) | 7.53 (-8.07, 22.77) |
| Age (MA) | | | | | | | | -6.86 (-22.70, 9.35) |
| Loo | 3361.2 | 3235.8 | 3138.2 | 3089.6 | 3090.3 | 3086.4 | 3086.2 | 3086.0 |
| ICC | 0.175 | 0.281 | 0.371 | 0.419 | 0.419 | 0.409 | 0.399 | 0.390 |
